# Do we understand orientation selectivity? A simulation study

**DOI:** 10.64898/2026.07.14.732589

**Authors:** Hugo Dictus, Alexis Arnaudon, Darshan Mandge, Joni Herttuainen, Armando Romani, Henry Markram

## Abstract

Orientation selectivity in the visual cortex has been extensively studied since it was disovered in 1959. In this article we demonstrate that current mechanistic explanations of orientation selectivity in the visual cortex, which appear to work in simplified models, do not suffice when embedded in a more realistic network. Up to now, orientation selectivity has only been reproduced in models with distance-dependent connectivity and limited cellular diversity. We therefore constructed a mouse primary visual cortex model with, among other refinements, realistic neuronal diversity and morphologically constrained connectivity. We implemented proposed mechanisms of orientation selectivity - specifically the organization of thalamocortical input into elliptical subfields and peferential connectivity between neurons with similar receptive fields - within this model. In the process we explored limitations of pure anatomy in predicting the connectivity and function of the cortex. Simulations of our model did not reproduce realistic levels of orientation selectivy. This indicates that these mechanisms, despite working well in simpler models, are not sufficient to explain orientation selectivity in real brain networks. On the basis of our results, we suggest the hypothesis that homeostatic plasticity is necessary to reconcile orientation selectivity with realistic connectivity.

## 1 Introduction

Many neurons in the visual cortex tend to respond more strongly to contours of some orientations than of others. This phenomenon, discovered in 1959 (Hubel & Wiesel, 1959) is called orientation selectivity. It has been extensively studied, and a large body of interature sheds light its mechanisms and role in visual processing.

To be accepted as fact, any explanation for this phenomenon must be demonstrably consistent with all of the relevant experimental data. In neuroscience this is challenging, as a very wide range of observations across a wide range of spatial scales must be incorporated simultaneously in order to demonstrate consistency with them. Through detailed and large-scale simulations, simulation neuroscience (Markram et al., 2011) offers a means of doing so.

Recent work on visual cortex models has shown how the extensive data available for this region can be integrated to construct empirically constrained explanations for key phenomena such as orientation and direction selectivity (Antolík et al., 2024; Billeh et al., 2020; Ito et al., 2026). Detailed models such as these allow us to demonstrate that a given explanation for a neural phenomenon is consistent with a wide range of anatomical and physiological data. The more such constraints a model incorporates the more realistic it is, and the greater the confidence we can have in it.

Models of the somatosensory cortex have achieved a higher degree of anatomical realism than those of the visual cortex (Isbister et al., 2025; Markram et al., 2015; Reimann, Bolanõs-Puchet, et al., 2024), incorporating detailed morphologies and statistically reconstructed synaptic connectivity. In particular, the connectivity of these models is more realistic than that of simpler models in terms of degree distributions (Giacopelli et al., 2021), and the expression of high-dimensional simplices (Reimann, Nolte, et al., 2017; Reimann, Santander, et al., 2024; Santander et al., 2025). These network features promote correlated activity, with strong and robust responses from a subset of neurons (Ecker et al., 2024; Santander et al., 2025), as observed in a variety of experiments (Barth & Poulet, 2012; Buzsáki & Mizuseki, 2014). However, these high-complexity approaches have not yet been applied to the visual cortex.

We constructed a detailed mouse primary visual cortex (VISp) model combining the structural precision of previous somatosensory cortex models with the functional features of previous visual cortex models. In the process we had the opportunity to test several hypotheses regarding how the anatomical features of the visual cortex shape its functional properties.

We aimed to incorporate two key features of the visual cortex, which were demonstrated in previous models to produce realistic orientation selectivity: orientation-selective thalamo-cortical input and like-to-like cortical connectivity.

The organization of thalamic inputs to cortical neurons is thought to shape their selectivity to orientation. The presence of ON (light responding) and OFF (dark responding) sub-regions of neural receptive fields have been observed in a variety of studies (Jones & Palmer, 1987; Martinez et al., 2005; Niell & Stryker, 2008; Ringach, 2004). In the mouse, such sub fields have been shown to be present in the sensory input from the dorsal lateral geniculate nucleus (LGd) and produce an orientation-dependent frequency modulation (F1-OSI) in the input current (Lien & Scanziani, 2013) in response to drifting grating stimuli. This is thought to be due to the synchronization of ON and OFF domains at the optimal stimulus orientation (Figure 5a). Differences in the temporal dynamics of sub-fields have in turn been demonstrated to produce direction-selective currents (Lien & Scanziani, 2018). Other visual cortex models used such an innervation scheme in combination with local like-to-like connectivity to reproduce orientation tuning (Antolík et al., 2024; Arkhipov et al., 2018; Billeh et al., 2020). Hebbian plasticity and waves of spontaneous activity are believed to shape this receptive field structure in-vivo (Bienenstock et al., 1982; Firth et al., 2005; Ge et al., 2021; Hunt et al., 2012; Martini et al., 2021).

Recent research found that ON and OFF thalamocortical boutons cluster together in the cortex (Tring & Ringach, 2022) and overlap with the spatial receptive fields of cortical neurons (Tring et al., 2022). This clustering has been suggested to be a consequence of the mosaic organization of the retina into ON/OFF dipoles (Paik & Ringach, 2011; Ringach, 2011). The authors proposed that this could generate an initial level of orientation selectivity which is later strengthened by plasticity. We implemented such a segregation and demonstrated that it produces negligible orientation tuning. We proceeded with theoretically optimal receptive fields as in Billeh et al., 2020.

Neurons in VISp preferentially connect to neurons with similar orientation preferences. Excitatory connections are more frequent between neurons sharing orientation preferences (Cossell et al., 2015; Ko et al., 2011; W.-C. A. Lee et al., 2016; Wertz et al., 2015), and connections between similarly tuned neurons tend to be stronger in both excitatory (Cossell et al., 2015) and inhibitory connections (Znamenskiy et al., 2024). Inhibitory connection probability could be influenced by like-to-like connectivity as well, but this is hard to assess due to the high density of inhibitory innervation in general (Bock et al., 2011; Fino & Yuste, 2011; Packer & Yuste, 2011). These trends are considered important in enhancing orientation selectivity (Antolík et al., 2024; Atallah et al., 2012; Billeh et al., 2020; S.-H. Lee et al., 2014; S.-H. Lee et al., 2012; Wilson et al., 2012) and producing orientation-tuned surround modulation (Adesnik et al., 2012; Angelucci et al., 2017).

Both receptive field overlap and response correlation are stronger predictors of connection strength than orientation preferences (Cossell et al., 2015), indicating that orientation-dependent connectivity trends are a consequence of general-purpose Hebbian plasticity mechanisms. We therefore applied a general connectivity rule based on the correlation of thalamocortical receptive fields, and found that this largely reproduced orientation-dependent connection probability.

Incorporating these two features into a detailed primary visual cortex model, we ran simulations and showed that orientation selectivity is lower than in-vivo. This contrasts with previous modeling results, suggesting that the added realism of the model somehow undermines its ability to reproduce this phenomenon. However, as it was not possible to comprehensively compare every aspect of this and previous models to experimental data, we cannot wholly rule out that the reduced orientation selectivity arises from some way in which our model is less realistic. If our interperetation is correct, it has implications for the interpretation of simpler models, calling into question the sufficiency of the mechanisms implemented. We suspect, on the basis of our results, that simulating realistic visual responses with realistic network anatomy requires the inclusion of homeostatic plasticity. Further analysis provides evidence consistent with this hypothesis. We argue more broadly that an account of orientation selectivity must follow from proposed plasticity mechanisms rather than static circuit properties in order to account for the true complexity of cortical connectivity and responses.

## 2 Results

### 2.1 Model anatomy

We briefly summarize the anatomical basis of the model, as all subsequent results build on it.

The model uses volumetric data derived from the blue brain atlas (Bolaños-Puchet et al., 2024; Erö et al., 2018; Piluso et al., 2025; Rodarie et al., 2021), which uses the Allen Institute common coordinate framework version 3. For the purposes of this study it was enhanced with geometric information on cortical orientation, depth, and layer boundaries (see methods for details).

We reconstructed only one hemisphere of the primary visual cortex, as this is all that was necessary for the phenomena we aimed to explore. We constructed depthwise cell density profiles for different excitatory and inhibitory types based on Schneider-Mizell et al., 2023 and Schüz and Palm, 1989. Neurons were assigned an mtype (morphological type) and etype (electrical type) as in previous work (Reimann, Bolanõs-Puchet, et al., 2024). They were additionally assigned a marker class (mclass) from EXC, Lamp5, Vip, Sst, and Pvalb, corresponding to excitatory, Lamp5-expressing, vasointestinal peptide-expressing, somatostatin-expressing, and parvalbumin-expressing neurons respectively. See methods 4.1.1 for details.

Neuronal morphologies for each neuron were synthesized based on the local cortical geometry as described in Kanari et al., 2022, while axons for them were selected from a pool of exemplars according to placement rules as in Reimann, Bolanõs-Puchet, et al., 2024.

We reconstructed the connectivity between neurons using previously developed algorithms (Reimann et al., 2015), applying anatomical constraints to the connectivity between neurons based on axo-dendritic touches (Figure 1d. Synaptic parameters were identical to those used for the somatosensory cortex.

**Figure 1:**
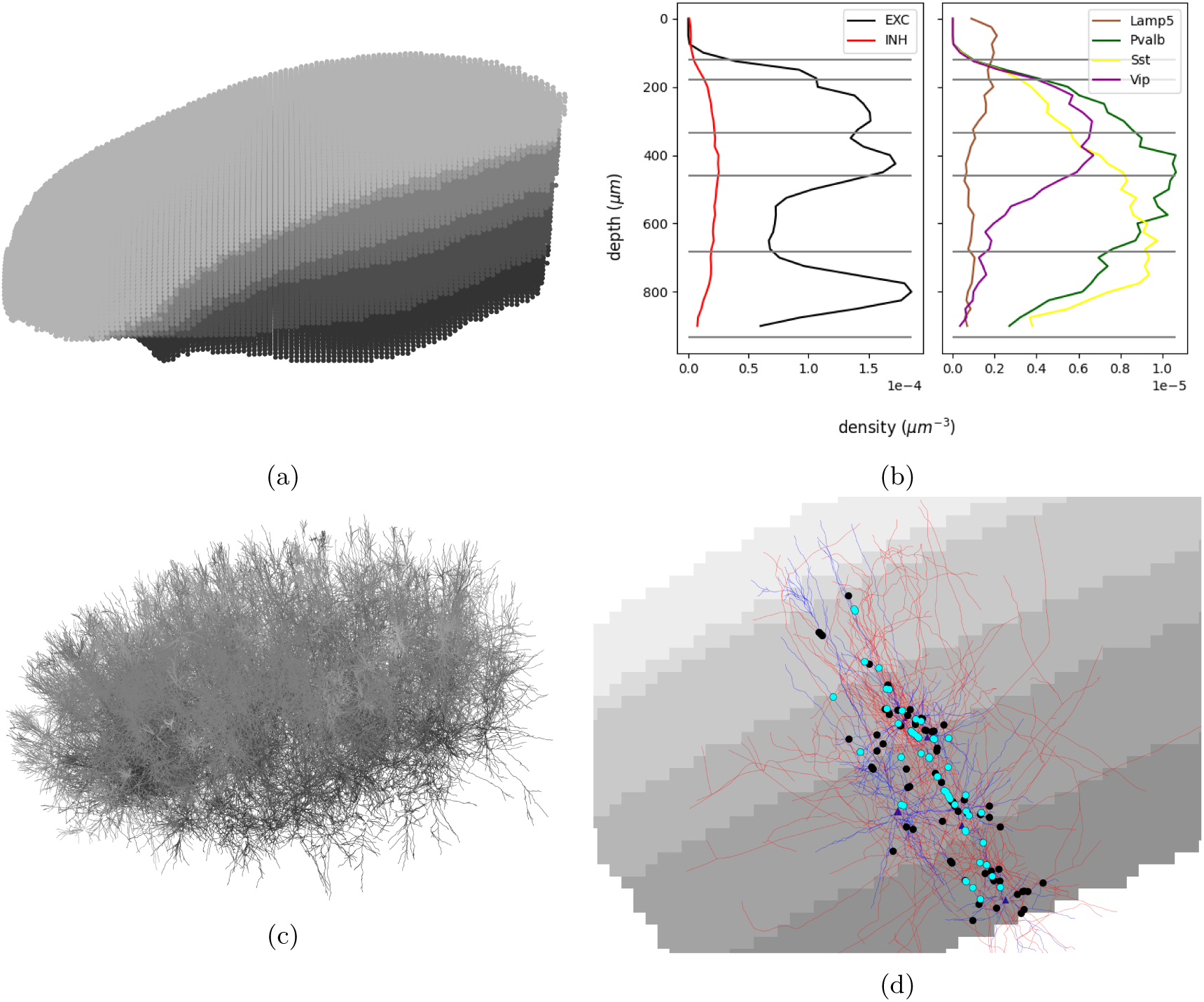
**a**: Geometry of the cortex according to the CCFv3 atlas. Shades of gray indicate cortical layers. **b**: Neuron density by depth. Gray lines indicate mean layer boundaries. **c**: Dendrites of 0.5% of the model neurons visualized. Shades of gray indicate soma layer. **d**: A subnetwork of four neurons, with connectivity based on anatomical constraints. Unused possible synapse sites are indicated by black circles, synapses by cyan circles

**Figure 2:**
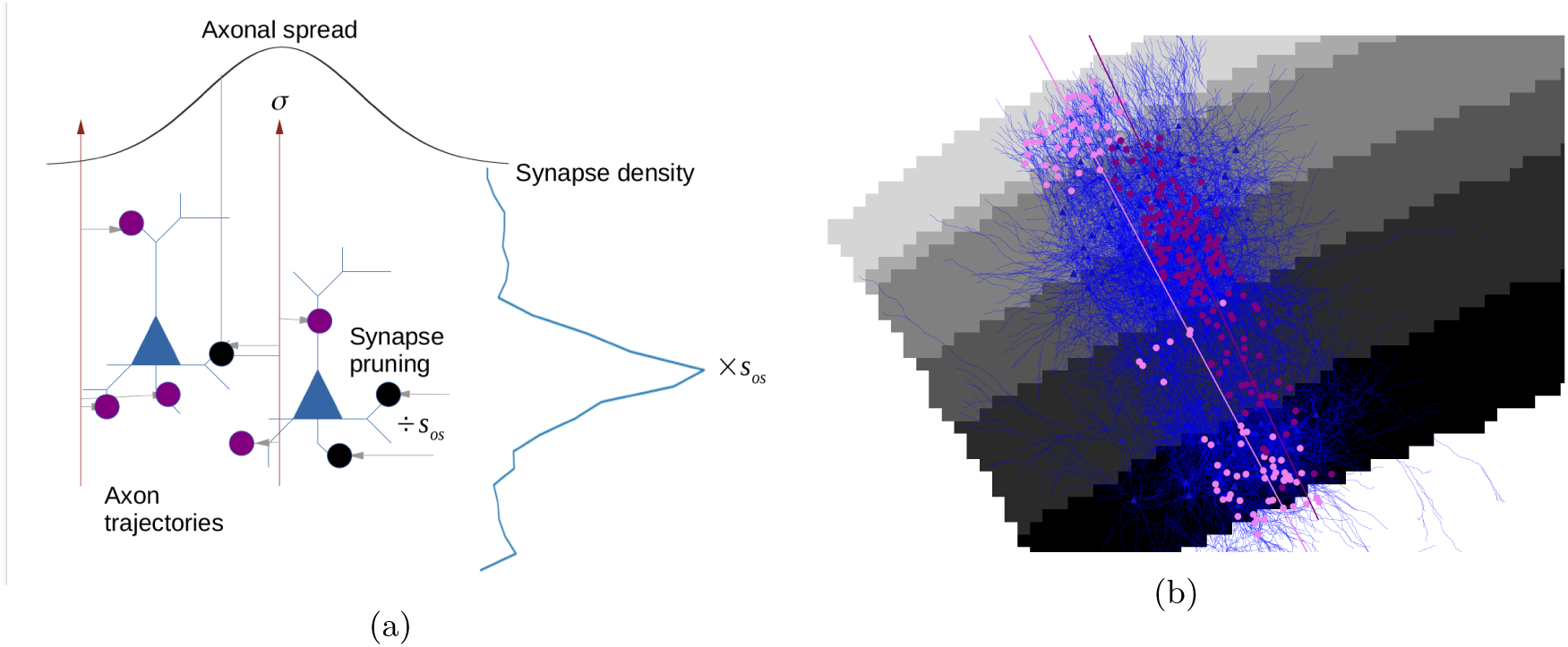
**a**: Diagram explaining the algorithm. Synapses are placed on dendrites according to a depthwise synapse-density profile, and oversampled by a factor *s_os_*. Subsequently they are assigned to presynaptic fibers with a probability proportional to a gaussian function of the horizontal distance between them with standard deviation *σ*. Finally, weaker connections are removed such that 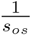 of the sampled synapses remain. **b**: Thalamocortical synapses from two exemplar LGd inputs, purple from the core and pink from the shell

**Figure 3:**
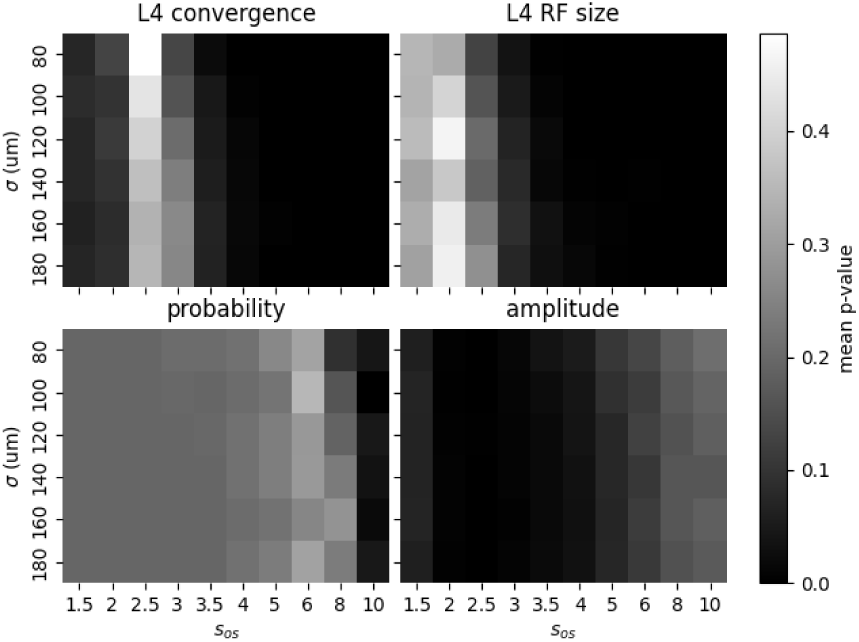
Scores (mean p-values) for different parameter combinations for each of the experiments. Scores shown are calculated for excitatory cell groups only. Note that the moderate score for innervation probability at low *s_os_* is mostly due to the p-value of 1.0 for predicting full innervation of layers 4 and 5. At these parameter values, complete innervation is predicted in all layers. The convergence and receptive field experiments can only be explained at fairly low oversampling, while innervation probability can only be moderately well explained at high oversampling and relative innervation amplitude is poorly explained by any parameterization. Extremely high oversampling values may lend themselves better to predicting relative innervation amplitude.

A retinotopic map spanning the visual field was constructed by mapping the span of the visual field reported in (Hübener, 2003) onto a flatmapped version of the atlas from (Bolaños-Puchet et al., 2024).

This anatomical scaffold provides a biologically grounded starting point from which we implement orientation selective receptive field structure and like-to-like connectivity.

### 2.2 Thalamo-cortical anatomy did not predict laminar and cell-type innervation patterns

To reconstruct thalamocortical connectivity, we used an algorithm developed previously for the somatosensory cortex (Markram et al., 2015; Reimann, Bolanõs-Puchet, et al., 2024). This algorithm uses the spatial distributions of dendrites and synapses to predict the thalamocortical input. We hypothesized that laminar and perhaps cell-type specific patterns of thalamocortical innervation could be reproduced by axo-dendritic overlaps as implemented by the algorithm. If true, an appropriate parameterization of the algorithm would reproduce experimentally observed patterns of innervation.

To test this, we varied the input parameters of the algorithm and compared the results to experiments. We found that both laminar and cell-type specific differences in innervation probability and strength were not accurately reproduced by any parameter combination.

The parameters *σ* (representing the horizontal spread of axons) and oversampling *s_os_* (affecting the number of synapses per connection) are not directly constrained by experimental data. We therefore varied these parameters to find the combinations that best explain (1) fraction of cells innervated by layer (Ji et al., 2016) (2) relative innervation strength by layer (Ji et al., 2016) (3) fraction of total thalamocortical excitation contributed by single connections in layer 4 pyramidal cells and (Lien & Scanziani, 2018) (4) receptive field size of layer 4 pyramidal cells (Niell & Stryker, 2008).

To evaluate the quality of each parameter combination we used the mean of p-values of tests comparing the experiment to the model. These can be thought of as representing the integral over model likelihood over the range of values more extreme than that observed from the experiment. The statistical tests used were:

1. binomial test for (1)
2. Mann-Whitney U-test for (2) and (4)
3. bootstrapping (see methods) for (3)

We used the mean of p-values as a metric rather than their product because every parameter combination had at least one p-value of 0, which would have rendered the score 0 for all parameter combinations had we used the product. Likewise, model likelihood was zero for all models, preventing its use as a metric.

Despite optimization, no parameter combination reproduced either laminar or cell-type differences in innervation. Best-fit parameters (*σ* = 100*µm*, oversampling=2.5) matched experiment 3 (fraction of excitation per connection) and 4 (receptive field size) well (p-values 0.44 and 0.161 respectively), but failed to reproduce laminar and cell-type specific targeting in experiments 1 and 2 (Figure 4, minimum p-values below floating point precision for both).

**Figure 4:**
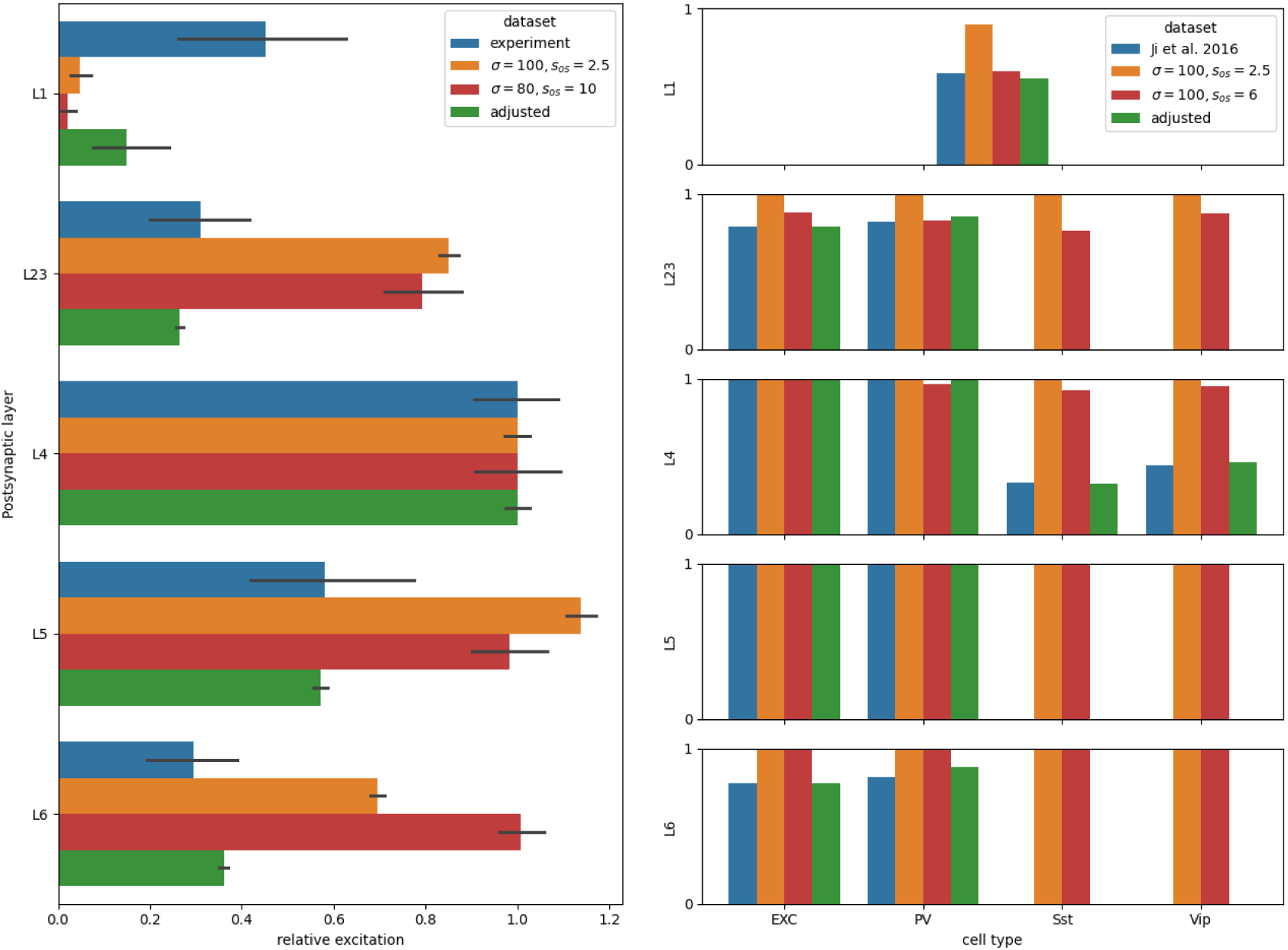
Comparison of three model parameterizations to (Ji et al., 2016), relative excitation strength by layer on the left and innervation probability on the right. In red is the parameter combination which best reproduced each particular result, but was not used. In orange the parameter combination with the best overall score across all four results Green is the version used going forward; it is the connectivity from orange after the it was adjusted to align with the experiments.

No parameter combination explained reduced innervation to Sst or Vip expressing neurons (Figure 4a). (Ji et al., 2016; Kloc & Maffei, 2014) (Figure 4b). Innervation probability in L1, L23, and L4 could be accurately predicted by some parameter combinations, although these parameterizations were unrealistic with respect to experiments 3 and 4. The innervation probability of layer 6 could only be predicted by parameter combinations which under-estimated innervation in other layers. Parameter combinations with adequate performance on 3 and 4 all predicted near-complete innervation of all layers, contrary to experiment. Likewise, the relative strength of innervation to different layers was not explained by any parameter combination (Figure 4b). The failure of anatomical factors to explain patterns in thalamocortical innervation indicates that other factors play a prominent role in them. However, we cannot rule out that a different algorithm, operating under different assumptions, could better explain the connectivity trends. We discuss this possibility in section 3.1.

We selected the parameter combination with the best mean score to adjust for succeeding modeling steps. As no parameter combination had any explanatory power for cell-type differences in innervation we excluded non-excitatory cells populations in calculating scores for experiments 1 and 2. We arrived at a result of 100*µm* spread and 2.5 oversampling. Our estimate of axonal spread is similar to that estimated previously from sparse axonal reconstructions (Reimann, Bolaños-Puchet, et al., 2022), which was 120*µm*.

Before proceeding with the rest of the study we corrected the remaining deficiencies to ensure the model has realistic innervation probability and strength for each cell type. Pruning synapses from over-innervated cell types to precisely match innervation probability from experiment 1 led to exceedingly weak innervation strength in layers 1, 23 and 6. Pruning the weakest connections to these cell types resulted in better layerwise innervation strength estimates, so we used this as our correction measure. Subsequently, we scaled synaptic strengths to each cell type to precisely match the mean relative innervation strength in experiment 2.

This corrected thalamo-cortical model was used for subsequent work.

### 2.3 Segregation of ON and OFF inputs to cortex did not reproduce thalamocortical orientation selectivity

Thalamocortical input to cortical neurons provides an initial level of orientation tuning through spatially offset subregions of the receptive field responding to increases (ON) and decreases (OFF) in illumination (Lien & Scanziani, 2013). With drifting grating stimulation, this produces orientation selectivity in the degree of frequency modulation in the thalamic input current (F1-OSI). It has been suggested that this could be anatomically implemented though the segregation of ON and OFF thalamic inputs (Tring & Ringach, 2022). We therefore hypothesized that by implementing ON/OFF dipoles in the thalamocortical connectivity of the model and simulating thalamic responses to drifting gratings we would reproduce substantial F1-OSI.

We constructed a map of ON and OFF centers across the surface of the cortex with Poisson-disc sampling (Methods). This map combined with the thalamocortical connectivity created a variety of receptive fields, both with and without clear ON and OFF subregions (Figure 5b).

**Figure 5:**
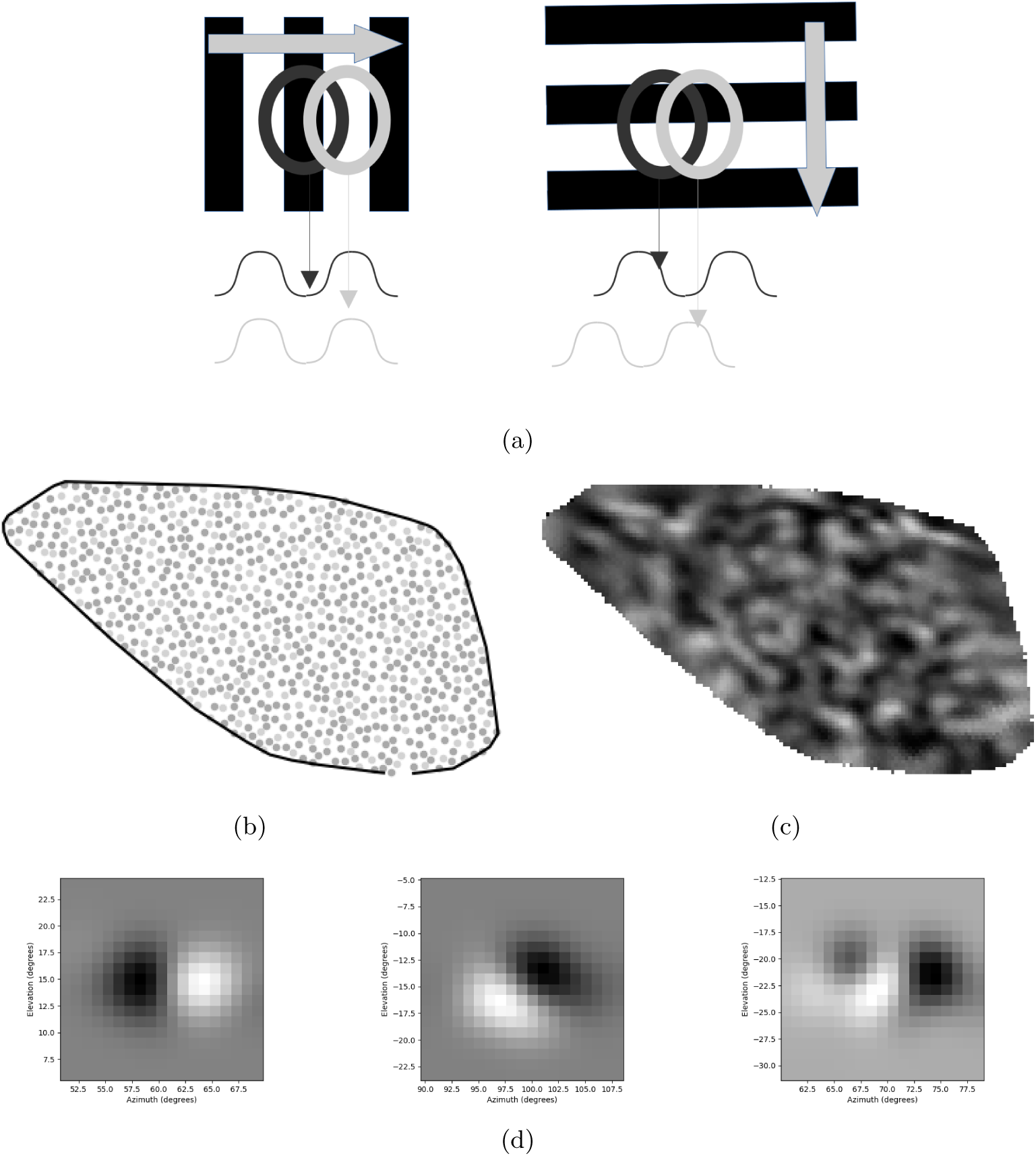
**a**: The mechanism for generation of orientation selectivity as observed by (Lien & Scanziani, 2013), proposed by (Tring & Ringach, 2022) to allow retinal ON and OFF domains to explain orientation selectivity. When a grating is presented at the optimal orientation the ON and OFF receptive fields synchronise their activation, leading to synaptic currents strongly modulated at the stimulus frequency. When the stimulus is non-optimal, the ON and OFF subregions are desynchronized, leading to a flatter overall input signal. **b**: visualization of ON/OFF cluster centers imposed on the flat coordinates of the visual cortex. **c**: Map of ON and OFF thalamocortical synapse density **d**: Receptive fields generated by anatomical segregation of ON and OFF input.

Using the brain-modeling toolkit (Dai et al., 2020) we simulated the responses of LGd neurons to drifting gratings at various orientations (see methods for details) and calculated the F1-OSI of the inputs to each cortical neuron based on spike histograms (Methods). Contrary to expectation, the model does not reproduce the experimental F1-OSI, and shows a similar distribution to a control model without segregation (Figure 6a, p=0.07, Mann-Whitney U-test), even when stimulus temporal frequency was varied S1). This indicates that the effect of ON-OFF input segregation in determining the orientation preferences of neurons is little to none.

**Figure 6:**
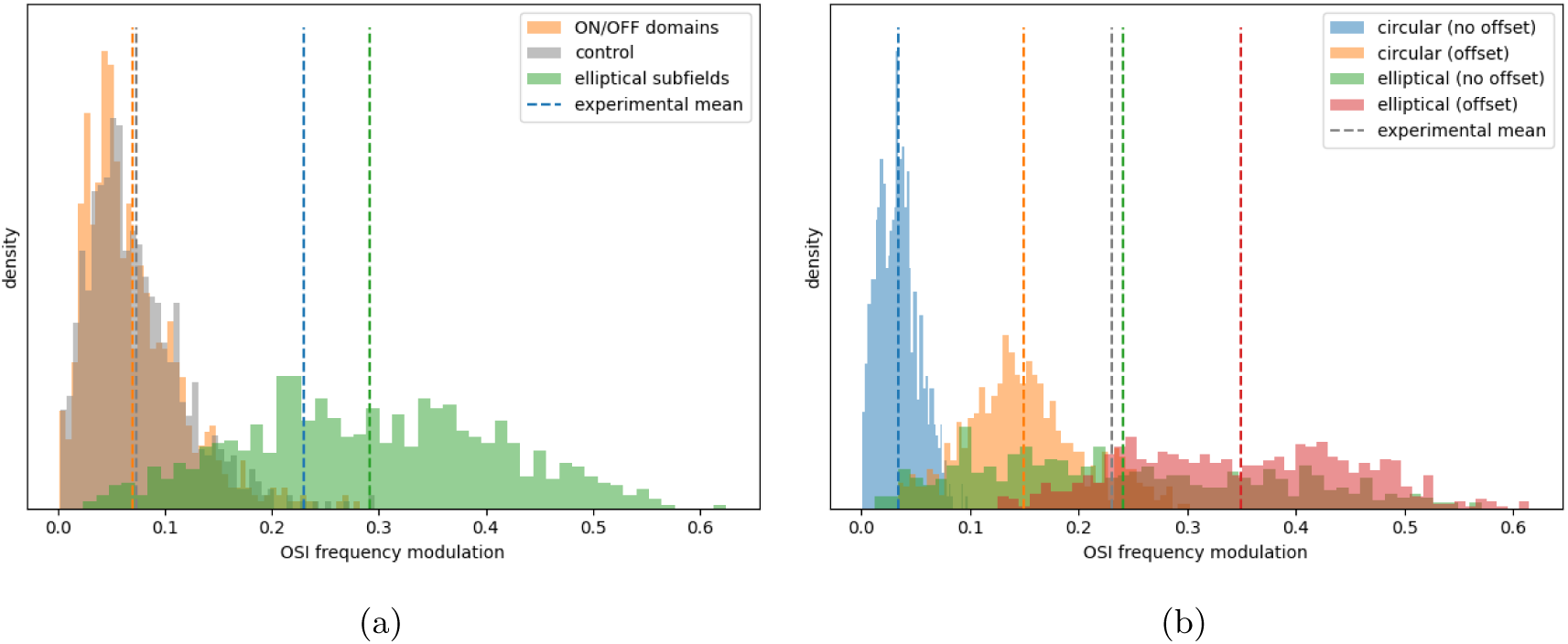
**a**: distributions of F1-OSI of input to layer 4 PCs for different model types, including a control model with no ON/OFF domains or aspect ratio, a model based closely on (Billeh et al., 2020) (elliptical subfields), and the model resulting anatomically from the introduction of ON/OFF domains (ON/OFF domains). **b**: F1-OSI for different receptive field models, circular ON/OFF subfields with no offset, circular with 10 degree offset, elliptical with no offset, and elliptical with 10 degree offset. Note that in these cases, sustained on and transient off subfields are used.

**Figure 7:**
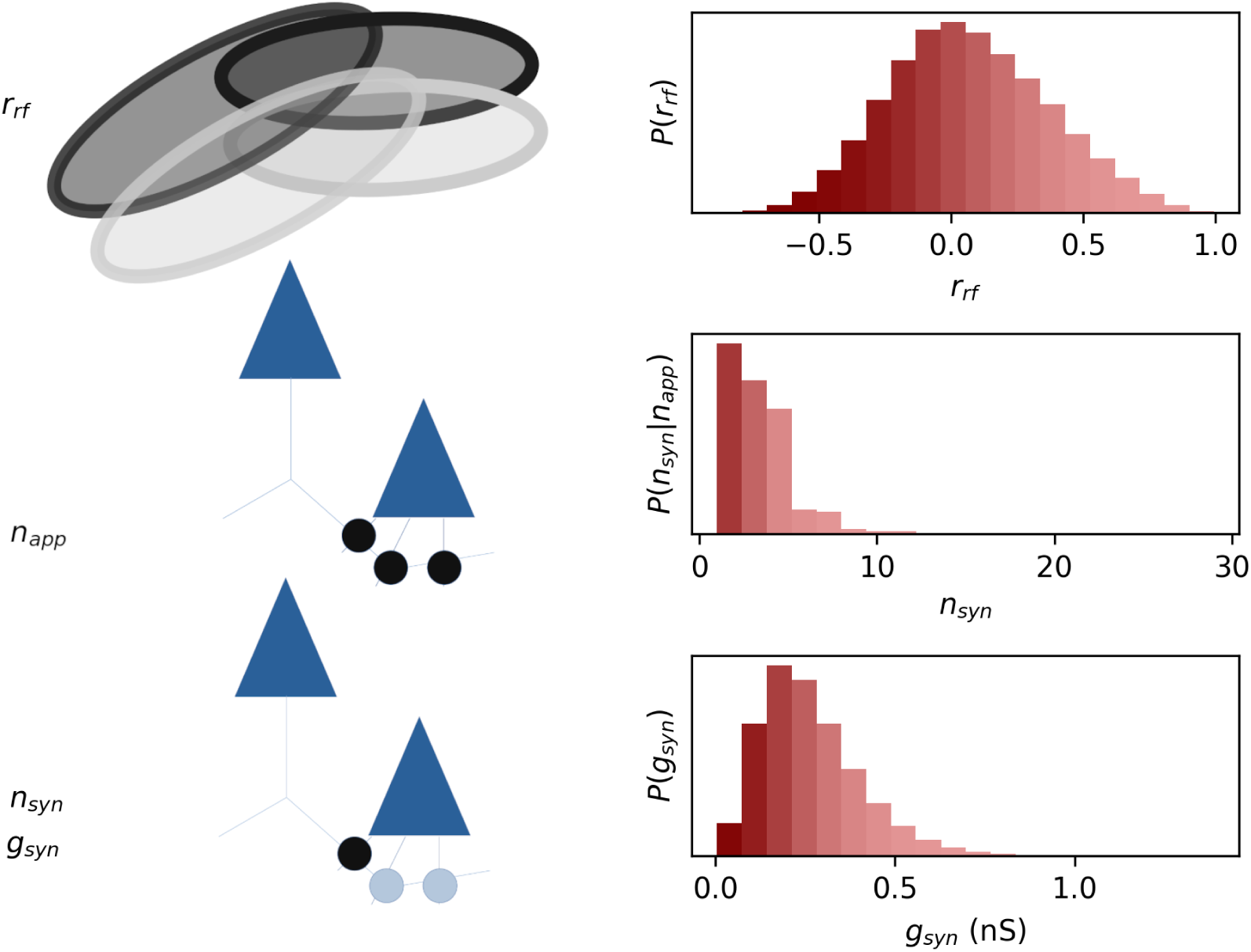
Diagram representing rewiring algorithm. The Pearson correlation *r_corr_* of the receptive fields of the neurons is calculated. From the *n_app_* potential synapse sites between a pair of neurons, *n_syn_* are instantiated for the lowest value of *n_syn_* where the cumulative probability of *n_syn_*is greater or equal to the cumulative probability of *r_corr_*. The synapse strengths *g_syn_* are re-distributed such that the synapse strength distribution similarly follows the *r_corr_* distribution. This maximizes the influence of *r_corr_* on connection probability and strength. The computation is run separately for each mclass-to-mclass pathway.

To understand why the experimental result was not reproduced despite the presence of ON/OFF offset we repeated the validation of F1-OSI while varying the aspect ratio and offset of the ON and OFF subfields.

Figure 6b shows that while both aspect ratio of the ON and OFF domains and their separation increase the level of modulation, the effect of aspect ratio is much larger and offset alone provides very little orientation tuning. However, the effect of clearly segregated ON and OFF subfields in these toy simulations is still significantly larger than the observed effects of ON/OFF domains in the circuit model. This could be due to the fact that the inputs to each neuron are strongest close to the center of its receptive field, as this is where the anatomical distance between axon and dendrite is statistically the lowest. In the toy models the strength of input is roughly equal at all points within the subfield. If we connected with equal probability and strength from all thalamocortical axons within a given radius, the offset of ON and OFF subfields may be more pronounced. We checked whether this produced any significant orientation tuning, and found it not to be the case (supplementary figure S8).

Our toy receptive field simulations revealed that orientation selectivity depends more strongly on the aspect ratio of ON/OFF domains than on their offset. Thus, the weak selectivity in the anatomically segregated model likely reflects the near-circular shape of receptive fields imposed by anatomy.

For all subsequent simulations we imposed elliptical sub-fields as in Billeh et al., 2020. See methods for the details of this conversion.

### 2.4 Maximizing like-to-like connectivity reproduced orientation and correlation-dependent connectivity

To reproduce the like-to-like connectivity observed in-vivo we reassigned synapses in the model based on the similarity of neurons’ receptive fields, while respecting the constraints of anatomy. The orientation and phase dependent rules of previous models intended to capture such relationships between neurons, and we here use those relationships directly. We analyzed the resulting connectivity to determine whether this approach reproduces the observed trends.

#### 2.4.1 Rewiring connections within anatomical constraints reproduced correlation-based connection strength

To generate the local connectivity in the model we used an algorithm (Reimann et al., 2015) previously used to reconstruct connectivity of the rat somatosensory cortex based on axo-dendritic contacts (Markram et al., 2015; Reimann, Bolanõs-Puchet, et al., 2024). This algorithm takes into account the anatomical features of the model, strongly constraining connectivity (Reimann, Horlemann, et al., 2017) and thereby reproducing connectivity patterns not explained by simple distance-dependent connectivity (Reimann, Nolte, et al., 2017; Reimann, Riihimäki, et al., 2022; Reimann, Santander, et al., 2024).

To impose like-to-like connectivity while maintaining the anatomy-connectivity relationship we implemented an approach that reassigns synapses in the model while respecting the conditional distribution *P* (*n_syn_* = *n*|*n_app_* = *k*) for each mclass-to-mclass pathway, where *n_syn_*is the number of synapses between a pair of neurons and *n_app_* is the number of appositions between them.

This approach starts by calculating the ON and OFF receptive fields of neurons in a grid and calculating the Pearson correlation coefficient of the receptive fields *r_rf_* for each pair of neurons. The correlation coefficients are then ranked and each pair of neurons assigned a quantile *q_rf_*representing the fraction of candidate connections with a lower *r_rf_*. They are assigned a number of synapses *n_syn_* based on this and the number of appositions *n_app_* such that *P* (*n_syn_ <*= *n*|*n_app_* = *k*) = *q_rf_*. This effectively maps the quantiles of the *r_rf_* distribution onto the *n_syn_*distribution. Synapses are then distributed and sorted by their weight *g_syn_* such that the strongest synapses are assigned to the most correlated pairs. See methods for details.

Importantly, this method maximizes the extent to which *r_rf_* determines connection probability and strength within the anatomical constraints. Cossell et al., 2015 analyzed the relationship of a number of variables to connection strength, including receptive field correlation and response correlation. There are two meaningful comparisons we can make to this experiment. Firstly, we can compare to the relationship observed with receptive field correlation to verify that our method reproduces its strength. Secondly, since we maximized the influence of *r_rf_* on connectivity we can compare to the strongest observed relationship, which was response correlation. The latter comparison allows us to assess whether the degree to which hebbian and anatomical factors determine the connectivity is realistic.

We hypothesized that this rewiring would greatly exaggerate the relationships observed in the experiment, as in the model the variable determining connectivity is precisely known and connectivity is maximally determined by it, while in the experiment response and receptive field correlation would have only been correlates of the variables which truly determined the connectivity, and measurements of all variables would be imperfect. Instead, the model matched the experiment surprisingly well for receptive field correlation (Figure 8a, compare orange and blue). However this figure does not account for the underlying distributions of the correlation variables, and therefore presents a stronger match than a quantitative validation. Half of connection strength was attributed to the 9% most correlated pairs of L23 neurons in the model compared to 12% in Cossell et al., 2015. The result of the experiment could feasibly have been sampled from the model (*p* ≈ 0.056, see methods).

**Figure 8:**
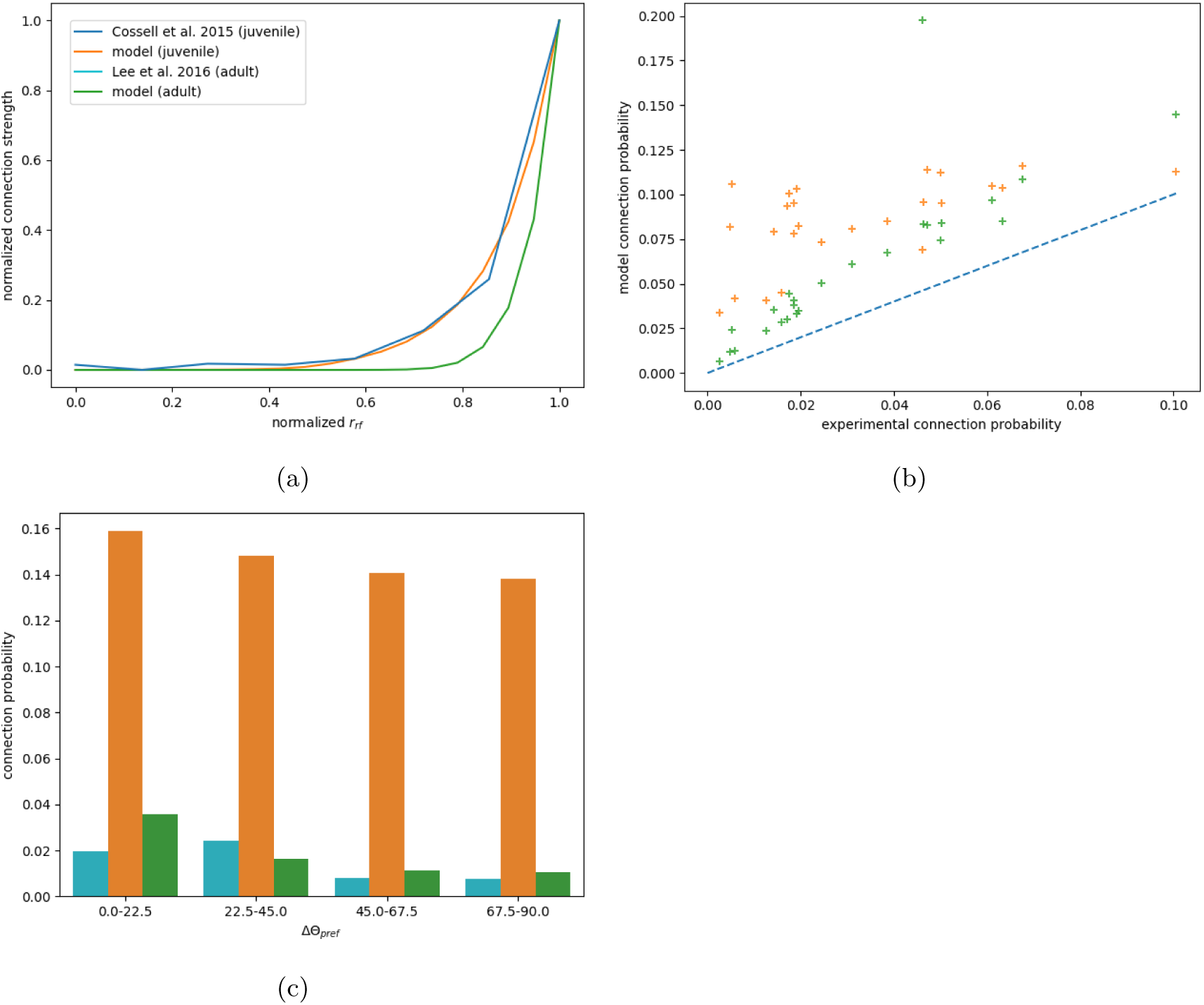
**a**: relationship between receptive field correlation and connection strength (including unconnected pairs) in the models and Cossell et al 2015, digitized from figure 2I. Both synapse strength and receptive field correlation are normalized so that the lowest value is 0, and the highest value is 1. Note that while the experiment calculated connection strengths from post-synaptic potentials, we used the sum of synaptic weights (maximum conductances *g_max_*) in each connection directly. **b**: Connection probability for original and adjusted model compared to MICrONs dataset, from cell type to cell type without layer distinctions. **c**: Connection probability by orientation preference difference compared between W.-C. A. Lee et al., 2016, the original model, and the recalibrated model

For response correlation, the experiment attributed 50% of connection strength to the 7% most correlated pairs, which is a closer match to the model (*p* ≈ 0.224, see methods). This is in spite of the fact that connection probability in the model is slightly sparser than in the experiment, and connection strength substantially more variable than in-vivo (see supplementary figures S4,S5). This could be taken to suggest that *in-vivo*, response correlation explains all variation in connection strength not explained by anatomical features, or else that the anatomical constraint provided by the model is too strong. However, various confounding factors prevent us from drawing either conclusion at present (see Discussion).

In short, the relationships produced in the model are plausible with respect to the data, either as a direct relationship between receptive field correlation and connectivity, or as a representation of the total hebbian component of connectivity.

### 2.5 Recalibration to adult-like connectivity reproduced orientation-dependent connection probability

This rewiring initially reproduced only a weak relationship between the orientation preferences of neurons and their connection probability (Figure 8c, orange). We considered that this was likely due to the substantially higher excitatory connection probability in our model than in this experiment.

Although the excitatory connection probability in the model is comparable to Cossell et al., 2015, which used juvenile mice, data sourced from adults shows substantially lower excitatory connection probabilities (Consortium et al., 2021; Jiang et al., 2015; W.-C. A. Lee et al., 2016). We additionally found that our connectivity algorithm struggled to accurately predict inhibitory connection probabilities from Schneider-Mizell et al., 2025.

In order to improve the realism of our model in these respects we re-ran the rewiring algorithm with a new estimate for each pathway of *P* (*n_syn_* = *n*|*n_app_* = *k*) derived from the MICrONs dataset (Consortium et al., 2021) (see methods for details). The result produced better connection probabilities (figure 8b), although it still overestimated connection probabilities overall (see Section 4.5.3).

Recalibrating the connection probabilities also led to a stronger relationship between *r_RF_*and connection strength than in the experiment (50% at 3%) (see Figure 8a, green). This is likely due to the lower connection probability in the modified model. The adjusted adult-like connectivity approximately reproduced the relationship between orientation preference and connection probability seen in W.-C. A. Lee et al., 2016 (Figure 8c), and the experimental result could plausibly have been sampled from the model (min. p-value=0.034 Bonferroni-corrected for 4 comparisons to 0.136).

We observed that although the connectivity was fit to layerless data, it predicted layerwise excitatory connectivity nearly equally well (Supplementary figure S6), while performing substantially worse for inhibitory connectivity (Supplementary figure S7). This difference is likely due to the absence of dendritic targeting rules for the inhibitory connections. This underscores the usefulness of circuit axo-dendritic anatomy for predicting layer-to-layer connection probabilities (Reimann et al., 2015), as well as the need to refine these for different inhibitory subtypes.

### 2.6 Orientation selectivity failed in the biophysical model

Up to this point we have only analyzed the patterns of input generated by the LGN model and the thalamocortical connectivity. We subsequently aimed to simulate the cortical network to reproduce orientation selectivity and other phenomena.

We initially ran simulations using biophysical neuron models in the NEURON simulator (Hines & Carnevale, 1997; King et al., 2009) as in Isbister et al., 2025, adapted using the methods from Arnaudon et al., 2023. These models had been validated with respect to their responses to step current injections as well as their dendritic integration properties, but not with respect to their threshold potential or responses to fluctuating current.

In these initial simulations, L4PCs showed orientation selectivity equal only to chance levels. We verified that both synaptic and somatic currents had realistic F1-OSI, and that the overall thalamocortical current and threshold currents were within a realistic range. When we injected the observed somatic current into simplified point-neuron models with parameters sampled from the Allen cell types database (Gouwens et al., 2019) we observed a small but significant degree of orientation tuning.

We suspected that the deficits in somatic integration in the biophysical models are due to their low threshold potentials, around or below −65 mV, which is substantially lower than seen in experiments (−30 to −55mV) (Baranyi et al., 1993; Contreras, 2004; Dégenètais et al., 2002; Gouwens et al., 2019; Williams & Stuart, 1999). This made them sensitive to brief, high-amplitude current fluctuations. We could not confirm this hypothesis, as we were unable to raise thresholds sufficiently by parameter tuning without abolishing spiking altogether.

Since point neurons demonstrated orientation selectivity while biophysical neurons did not, we converted the model to the NEST simulator (Gewaltig & Diesmann, 2007) to ensure that cell-intrinsic issues did not obscure network phenomena. See methods for the details of this conversion.

### 2.7 Calibration reproduced plausible firing rate distributions

Our model does not represent all of the inputs to the primary visual cortex, so to reproduce realistic firing rates we needed to provide additional excitatory input to compensate for the missing synapses. We used synapses without short-term plasticity firing at 1000Hz to represent input from many different afferents. We scaled the weights of these to a fraction of the threshold current of the target neuron, with different fractions for different layers and to Pvalb and non-Pvalb populations. We simulated thalamic responses to a gray screen stimulus and varied the strength of compensatory synapses to different cell types to reproduce a fixed fraction of the firing rates observed with a gray screen in Siegle et al., 2021 (see methods for details).

The target fraction of experimental firing rates was determined by comparing spontaneous firing rates of Pvalb-expressing neurons from W.-p. Ma et al., 2010 with those of fast-spiking neurons from Siegle et al., 2021. Because all Pvalb-expressing neurons in the experiment exhibited non-zero firing rates, it was unclear whether this ratio should include or exclude neurons that were silent in our simulations. To address this, we performed two calibrations: one including all neurons, and another considering only those active during the 10-second simulation. When silent neurons were excluded, thalamic input contributed 90% of the total excitatory current to L4PCs during drifting grating stimulation. In contrast, including silent neurons yielded a fraction of 34%, aligning with the one-third value reported in Lien and Scanziani, 2013. Notably, the extent to which excluding silent neurons inflates mean firing rates depends on simulation length, as shorter simulations tend to yield a higher fraction of silent neurons. For these reasons, we adopted the calibration that included all neurons in the firing rate calculation.

While our calibration undershot the targets (Figure 9b), the firing rates were nonetheless consistent with the cell-attached recordings for Pvalb-expressing neurons (Figure 9d). The firing rate distribution was strongly tailed (Figure 10c), which is consistent with experiments (Buzsáki & Mizuseki, 2014). The tailedness seen in the model may be somewhat exaggerated, as it has maximum firing rates double those observed in Siegle et al., 2021, and around a third of Pvalb-expressing neurons are silent, in contrast to W.-p. Ma et al., 2010. However, due to the small fraction of highly active neurons in the former case and the shorter duration of the simulation in the latter case, the model distribution cannot be ruled out by these data.

**Figure 9:**
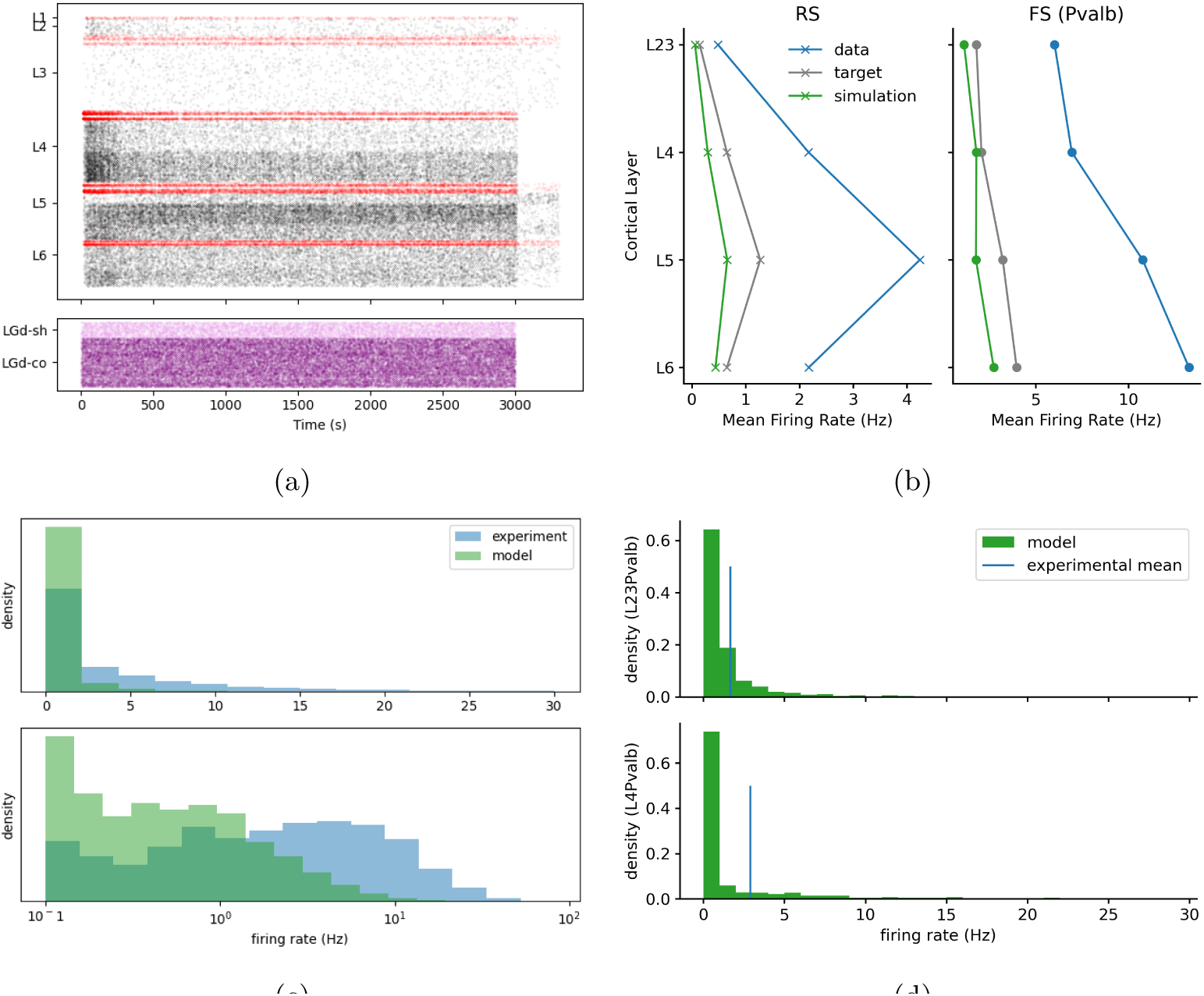
**a**: Raster of activity in gray-screen spontaneous activity. **b**: Firing rate per cell type compared to Siegle et al., 2021. Note that the target values were a fixed fraction (32%) of the experiment. **c**: Histogram of firing rates for excitatory neurons in the model (green) compared to RS neurons in Siegle et al., 2021 (blue), linear scale above, logarithmic below. **d**: Comparison of spontaneous firing rates for Pvalbumin-expressing neurons to W.-p. Ma et al., 2010. p-values from bootstrapping test were 0.165 and 0.206 for L4 and L23 respectively.

**Figure 10:**
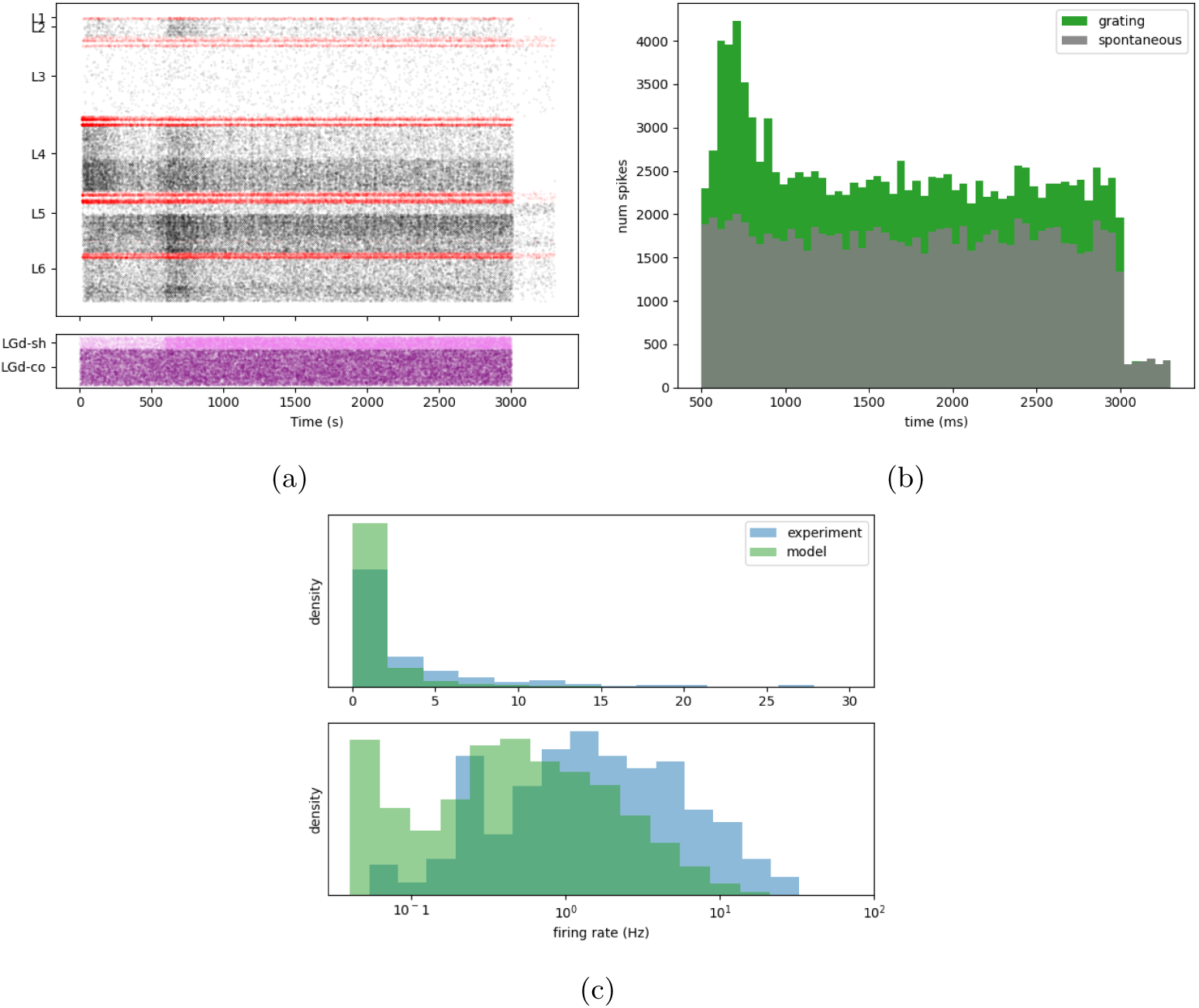
**a**: Raster of spiking activity in a drifting grating simulation with temporal frequency of 4Hz. **b**: PSTH compared between spontaneous activity and drifting grating stimulation. A transient peak in activity is observed, as in-vivo. **c**: Distribution of model maximum firing rates and logarithm therof (green) compared to Siegle et al., 2021 (blue)

A raster plot of the spontaneous activity in the calibrated model can be seen in figure 9a. An in-vivo like asynchronous state can be seen, as expected.

We also simulated responses to drifting grating stimuli (Figures 10a and 10b). There is a clear transient peak in activity at stimulus arrival, and a smaller, sustained increase in activity afterwards. Maximum firing rates (averaged over trials) per neuron are plausible with respect to Siegle et al., 2021 (Figure 10c), considering its sampling biases.

### 2.8 Model orientation selectivity was significantly lower than experiment

Our model implements orientation-dependent connectivity and orientation-selective receptive fields, which in previous models reproduced realistic levels of orientation selectivity (Antolík et al., 2024; Arkhipov et al., 2018; Billeh et al., 2020). We therefore hypothesized that it would likewise do so.

First we disabled all cortico-cortical synapses, with background input appropriately adjusted (supplementary figure S11), and simulated responses to drifting gratings. Orientation selectivity was distinguishable from control (p<0.005 for all layers and cell types, p<10*^−^*^99^ for L4PCs, Mann-Whitney U-test), showing that the mechanisms of orientation selectivity are working, but weaker than in Siegle et al., 2021 (lowest p < 5 × 10*^−^*^10^, Mann-Whitney u-test) (Figure 11a), showing that they are not sufficient.

**Figure 11:**
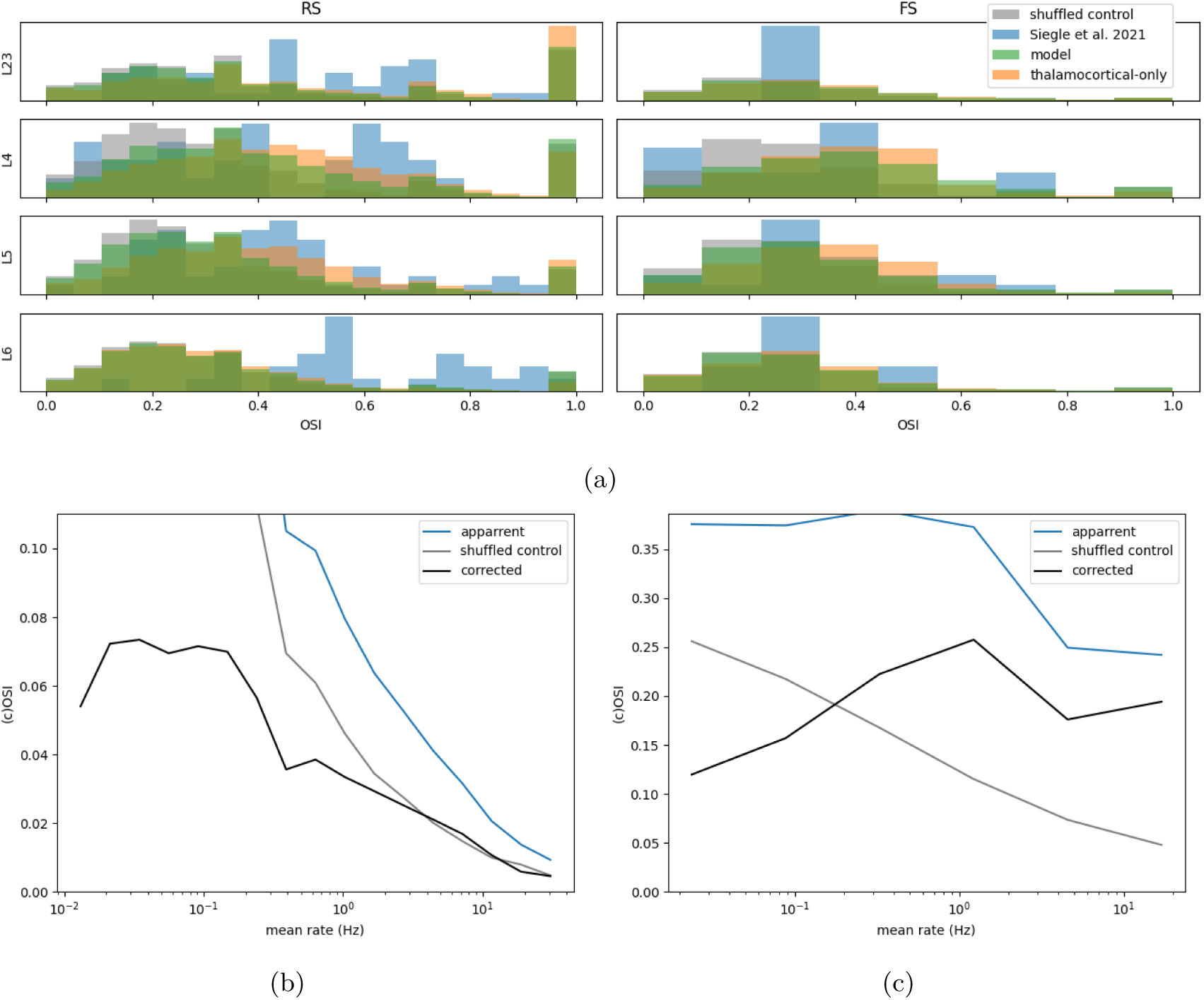
**a**: Orientation selectivity index compared to Siegle et al., 2021. Blue is the experiment, gray is a shuffled control model, orange the thalamocortical-only model and green the full model. **b**: Mean cOSI by mean firing rate for EXC neurons (black), with OSI for the model and statistical control in blue and gray. **c**: like b, but for Siegle et al., 2021

When we ran simulations with the full model we found that including the local connectivity reduced OSI rather than enhancing it (Figure 11a, compare orange and green). This result suggests that the incorporated mechanisms are not sufficient to produce realistic levels of orientation and direction tuning, which is in conflict with the results from previous models (Antolík et al., 2024; Billeh et al., 2020).

This raises the question: which of the differences between our model and previous work explains the different restult? The principal difference between our model and previous models is the more realistic anatomically-constrained connectivity. Given that in our model the addition of local connectivity disrupts orientation selectivity, rather than enhancing it as in Billeh et al., 2020, it seems likely that this is the source of the difference. The introduction of more realistic connectivity patterns may disrupt the efficacy of mechanisms that generate orientation selectivity in simpler models.

This would indicate that the implemented mechanisms are not, by themselves, sufficient to explain orientation selectivity in real brain networks. This then raises the additional question of which missing mechanisms are required for orientation selectivity in-vivo.

#### 2.8.1 The relationship between firing rate and OSI differs from in-vivo

To better understand why local connectivity did not enhance orientation selectivity we quantified the capacity of each neuron to impart an orientation-selective signal to its postsynaptic partners. We defined a control-corrected orientation selectivity index (cOSI) determined by subtracting the OSI of a shuffled control of the neuron’s responses from an OSI of its raw responses (in both cases without subtracting a baseline value), which should more accurately represent the extent to which a neuron’s outputs reflect orientation signals than conventional indices.

We found that cOSI was higher for neurons with lower firing rates and was dramatically reduced for neurons with high firing rates (Figure 11b). The same relationship was not observed in Siegle et al., 2021, where OSI was fairly independent of firing rate and control OSI reduces with firing rate, resulting in a net increase in cOSI with firing rate (Figure 11c).

We suspected on this basis that the lack of homeostatic balancing at the single-neuron level explains the reduced orientation tuning compared to experiment. We therefore implemented a per-neuron calibration procedure designed to match neuronal thresholds to their mean inputs, in hopes of mimicking the effects of homeostatic plasticity and thereby recapitulating orientation selectivity.

### 2.9 Network effects destabilized balanced input conductances

We assumed that for each neuron type there is a fixed ratio *f_g_* of threshold current *I_th_*to mean current at threshold potential *I_µ_*, s.t. *f_g_I_th_* = *I_µ_*. This can be expanded to an expression relating mean leak, excitatory, inhibitory, and background conductances *g_l_, g_e_, g_i_, g_b_* to the threshold and reversal potentials *V_t_h, V_e_, V_i_*, *g_l_*(*V_th_* − *V_l_*) = (*g_e_* +*g_b_*)(*V_e_* −*V_th_*)+*g_i_*(*V_i_* −*V_th_*) We used simulations with poisson inputs at the target rates to to predict *g_e_, g_i_* from connectivity and adjusted *V_th_*and *g_b_*on a per-neuron basis to satisfy the constraint for a given value *f_g_*. We varied *f_g_* with poisson inputs of to determine the correct value of *f_g_*for each mclass. See methods for details.

We expected that this would produce fairly even firing rates within each mclass. In the idealized simulations, a fairly realistic distribution of firing rates was observed (Figure 12a). However when we applied the method to a full simulation (Figure 12b) the skew and tailedness of the firing rate distribution were dramatically amplified.

**Figure 12:**
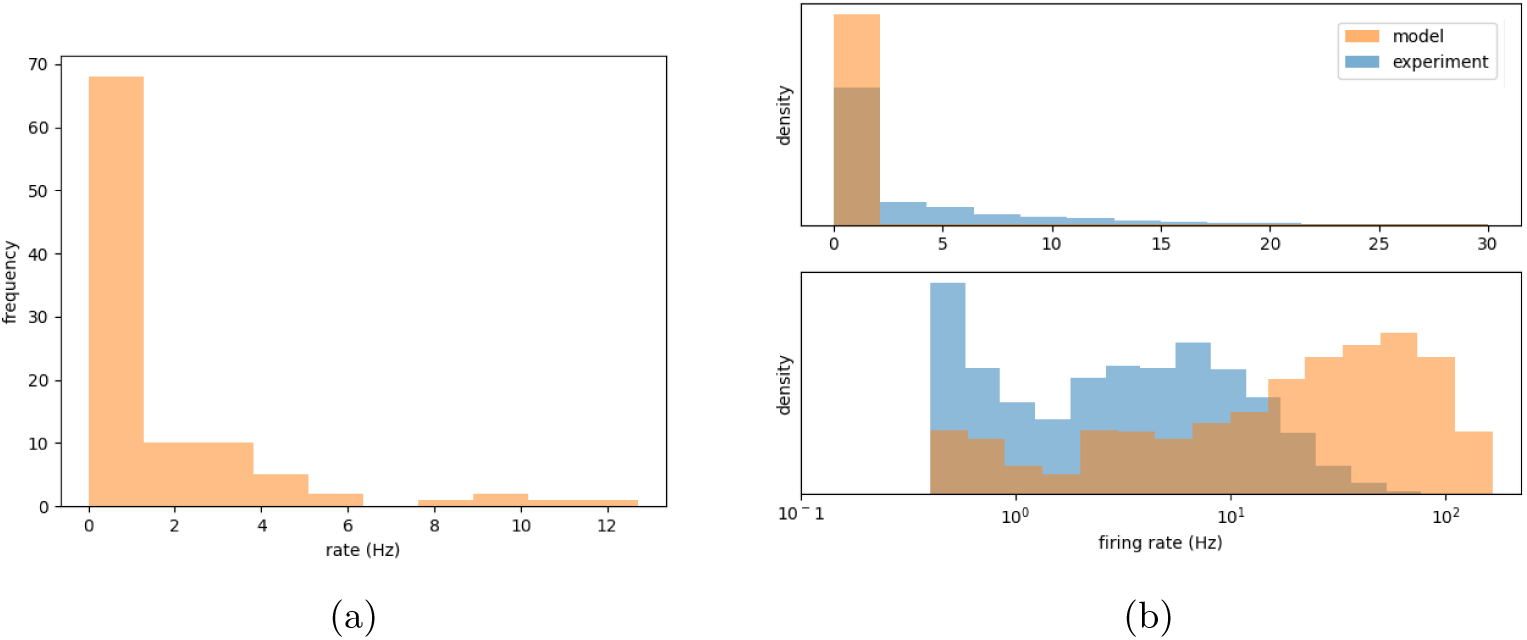
**a**: Firing rate of L5PCs with per-neuron input calibration in idealized simulations **b**: Firing rate of L5PCs with per-neuron input calibration in a real simulation (orange) compared to Siegle et al., 2021 in blue. Note that the tail of the former extends to 200Hz.

**Figure 13:**
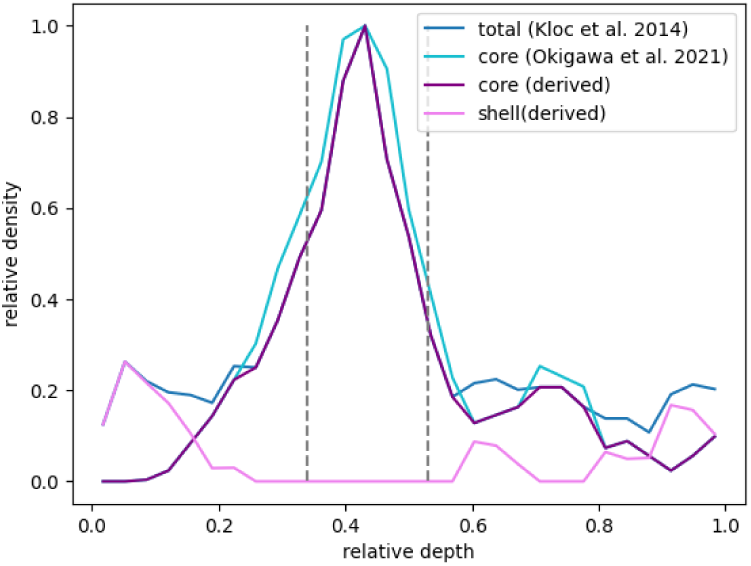
Measured and estimated relative axon density by depth for core and shell pathways. The shell and core densities were derived by lining up the peaks of the two datasets and subtracting core-only axon density from total axon density

The failure of our approach is in the generalization from the idealized case to the fully connected network. This suggests that imbalances in neural inputs and excitability are strongly influenced by network effects. We therefore suggest that simulating homeostatic plasticity online would be a more promising means of achieving network balance, but new methods would be required to permit simulation on the necessary timescales.

## 3 Discussion

The inability of our model to explain key features of the visual cortex illustrates, through their absence, the importance of dynamic plasticity processes in visual perception and cortical function. Significantly, it does so in ways which cannot be achieved with simpler modeling approaches, with which the necessity of plasticity is less obvious. Each of our results relates to this theme, and the relevant insight is only possible because of the high verisimilitude of the modeling approach. As others have demonstrated, phenomena such as orientation selectivity can be reproduced by computational models without plasticity, but we suggest that the success of this approach is conditional on the omission of realistic network features, limiting its utility for understanding cortical function in-vivo.

As our conclusions stem from the lack of explanatory power in our model, they must be received with some caution. Although we strove to maximize the realism of the model, we cannot rule out that the same data, incorporated in a different way, could produce more accurate predictions of the properties of the visual cortex. Neither can we rule out that inaccuracies or uncertainties in the experimental data mask the explanatory power of our model.

### 3.1 Does axon density determine thalamocortical connectivity?

Our algorithm failed to predict, on the basis of dendritic morphology, cell positioning, and axon-fluorescence data, the patterns of innervation probability and strength to different cell types and layers. This suggests that the targeting of thalamocortical synapses, while influenced by axonal projection density, is strongly affected by other factors, such as differential expression of connectivity-guiding molecules (Yamamoto & Hanamura, 2005) including Lhx2 (Wang et al., 2023), or synaptic competition from local and long-range axons. There are some sources of uncertainty in our analysis.

Firstly, our compositional mapping of the Schneider-Mizell et al., 2025 m-types to BlueBrain ME-types (Methods 4.1.1) is likely imperfect. Additionally, there may be minor differences in dendritic morphology between rat somatosensory cortex and mouse visual cortex. We could find no direct evidence of differences in dendritic morphology between rat and mouse neocortex besides overall size (which we accounted for), and no direct evidence of differences between primary visual and somatosensory cortices. Some variation between different regions has been observed in mouse (Benavides-Piccione et al., 2006), but primary sensory cortices were not compared. Studies of primate cortex suggest that primary sensory cortices have similar morphologies and inter-region variation is driven by differences between sensory and association cortices (Bianchi et al., 2013). On the basis of this we expect any inaccuracies introduced by our morphologies to be minor and insufficient to explain the failure of our model to reproduce layerwise innervation.

Our estimate of the number of thalamocortical afferents assumes that all thalamocortical projection neurons in LGd target the primary visual cortex. It additionally assumes a larger number of projection neurons than estimated in Evangelio et al., 2018. Our estimate is therefore an upper bound. A smaller number of thalamocortical afferents would have necessitated smaller oversampling and larger spread to explain the fraction of excitation per connection from Lien and Scanziani, 2013. At these low values of oversampling, layer-wise differences in innervation would be even less pronounced in the model, strengthening our conclusions.

Furthermore, the axonal projection patterns and validation data we used came from different studies and therefore different subjects. If there is large inter-subject variation in these properties then the failure of our model may not necessarily reflect a failure of its assumptions, undermining our conclusions.

Our estimate of peak thalamocortical synapse density was substantially lower than a recent estimate in the mouse barrel cortex Bopp et al., 2017. If the true synapse density in the visual cortex is comparably high, then innervation probabilities predicted by anatomy would be higher across the board, strengthening our conclusions.

In calculating the thalamocortical synapse density (Methods 4.2.1) we assumed that the synapse density is proportional to the axon density, and the axon density is proportional to the fluorescence observed in the studies used. Either of these assumptions could be mistaken.

It is possible that rather than being proportional to axon density, synapse density is greater where there is also a high density of dendrites. In this case the reduced innervation of layer 6 may be easier to explain in anatomical terms, while the reduced innervation of layers 1 and 2/3 would be more difficult to explain. As all parameter combinations which had realistic fractions of excitation per connection estimated complete innervation of all layers, we believe our conclusions would be unchanged in this case.

It is also possible that synapse density scales super-linearly with axon density with a different form of cooperative synapse formation. Our algorithm prunes every morphological type separately, removing the same fraction of synapses from each in order to preserve the shape of the synapse density distribution. If instead the same cutoff point was used for all cell types then cell types receiving fewer synapses overall would be disproportionately targeted by the pruning, and layer-wise differences may be more pronounced.

Finally, the most challenging of the limitations is the possibility that axon fluorescence scales sub-linearly with axon density, which is plausible due to the scattering of light in brain tissue. If the relationship is strongly sub-linear, then the axon density distribution would have sharper peaks and troughs than in our estimate, and may more accurately predict layer-wise innervation.

Under the assumption that the relationship between axon fluorescence and axon density is approximately linear, our results allow us to conclude that axo-dendritic anatomy does not straightforwardly predict layerwise innervation: synapse density is not directly proportional to axon density. Since the less-innervated layers tend to receive strong cortico-cortical inputs (Harris et al., 2019; Shen et al., 2022), it may be that some form of competitive plasticity is responsible for this non-linearity. It may also be that part or all of this nonlinearity is explained by cooperative synapse formation, leading to axons forming fewer synapses in layers where their density is lower. We conclude, conditioned on an approximately linear relationship between axon fluorescence and density, that thalamocortical synapse density does not scale linearly with thalamocortical axon density. We firmly conclude that anatomical factors do not explain cell-type differences in thalamocortical innervation, and suggest that molecular targeting mechanisms determine this in-vivo.

### 3.2 Do thalamocortical ON/OFF domains shape cortical orientation selectivity?

Our results show that imposing a segregation of ON and OFF inputs to the cortex does not generate substantial orientation tuning in layer 4 pyramidal neurons. This indicates that the effect of such segregation on orientation selectivity is negligible. Supporting this is the observation that in early spontaneous retinal activity, ON and OFF retinal ganglion cells fire in synchronous waves (Kerschensteiner & Wong, 2008; Wong et al., 1993), so that by the time ON and OFF segregation plays a role in cortical development initial orientation preferences may already have developed. The observations of Tring and Ringach, 2022 and Tring et al., 2022 may be better explained by plasticity mechanisms forming orientation maps in cortex (Miller, 1994; Najafian et al., 2022), rather than by segregated inputs.

Our results instead support the primacy of Hebbian plasticity in both thalamocortical receptive field formation and polar map formation in mouse VISp. However, the fact that our implementation of this mechanism did not generate F1-OSI does not mean that a modified version of it could not. In particular, a modified version in which transient and sustained inputs are also segregated, exerting a subtle influence on tuning, may still be tenable.

### 3.3 How strongly do anatomy and like-to-like trends determine connectivity?

As described in section 2.4.1, our model reproduced a relationship between receptive field correlation and connection strength of similar strength to that observed between response correlation and connection strength in Cossell et al., 2015. When recalibrated to match adult connection probabilities also approximately reproduced the relationship between orientation preferences and connectivity.

If we accept the anatomical constraint imposed by the model to be accurate, this result suggests that in the visual cortex of the juvenile mouse, response correlation is a remarkably strong predictor of connectivity and connection strength, explaining all or almost all variation within the constraints imposed by anatomy. Conversely, it may be the case that the anatomical constraint we impose is too strong. Unfortunately, a number of confounding factors prevent us from drawing strong conclusions either way. Firstly, it may be the case that in-vivo there is a relationship between the similarity of thalamocortical inputs to neurons and the number of appositions between them (but see W.-C. A. Lee et al., 2016), which is not accounted for by our model due to the loss of spatial information in the receptive field assignment step. Secondly, the presence of a strong connection between two neurons can cause them to fire in a more correlated fashion, as explored by Ecker et al., 2024. Thirdly, the experiment of Cossell et al., 2015 may have induced plasticity in the activated connections of correlated neurons, thereby amplifying the observed correlation-connectivity relationship. As these factors cannot be quantified at present, we cannot draw strong conclusions in this case.

The parameter values found in recalibrating the model connectivity (see methods section 4.5.1) support strong anatomical constraint on connectivity. We found the optimal value of the *a* parameter was positive for all pathways, predicting that pairs of neurons with more axodendritic touches will have a higher fraction of those touches converted to synapses on average. This is consistent with the principle of cooperative synapse formation (Fares & Stepanyants, 2009) which informs our reconstruction approach.

Both the original and recalibrated models overestimated the distance-dependence of connection probability, while the conditional probability distribution calculated during the calibration procedure predicted a more realistic profile (supplementary figures S10). It may be explained by the fact that neurons which are closer together are more likely to have high receptive field correlation and therefore are more likely to connect. This effect was not taken into account in the calibration, leading to an exaggerated distance dependence. If the distance-dependence of connectivity is partly explainable in terms of correlation, we may therefore be over-estimating the influence of anatomy on connection probability. Analysis of touch-synapse relationships in large-scale electron-microscopy datasets like Consortium et al., 2021 would allow the direct quantification of the relationship between anatomical touches and synaptic connectivity. Conditional synapse count distributions could then be used to directly initialize realistic connectivity-anatomy constraints, without the need for additional calibration.

Our results here suggest, subject to uncertainty, that plasticity in cortico-cortical synapses fully utilizes the degrees of freedom permitted by the anatomical constraints, with little to no noise in the determination of synaptic connectivity.

### 3.4 Orientation selectivity in the cortical network

Our model’s reduced orientation selectivity compared to experimental data and previous work prompts two questions. First, which in vivo mechanisms essential for orientation selectivity are missing from the model? Second, how might our model’s added complexity disrupt the selectivity observed in simpler models? Although we could not definitively answer these questions, we propose some possibilities for further research.

A significant difference between our model the previous ones is the more complex, anatomically constrained connectivity. This connectivity leads to more realistic, long-tailed degree distributions (Giacopelli et al., 2021) and higher-order network structure that promotes chaotic dynamics (Nolte et al., 2020; Santander et al., 2025), pushing networks to the "edge of chaos" where computational capacity is maximized (Bertschinger et al., 2004; Legenstein & Maass, 2007). We hypothesize that this refinement in our model over previous models is responsible for the difference in our results.

In-vivo, neurons are exquisitely balanced for robust and efficient representation through homoeostatic and other plasticity mechanisms (Davis & Bezprozvanny, 2001; Z. Ma et al., 2019; Pratt & Aizenman, 2007; Turrigiano, 2012) which may be important for orientation selectivity (Cogno & Mato, 2015). We suspect that this is the most important missing mechanism for orientation selectivity. Two pieces of evidence support this suspicion.

The first piece is the difference between our model and previous ones. The absence of homeostasis in our model is likely to be more impactful than in simpler models due to the long-tailed degree distributions and complex network structure. In simpler models, the amount of input and output to each neuron is partially balanced by the fairly narrow per-type degree distribution and limited higher-order structure. This is further evidenced by our result that the introduction of network effects drives divergence in firing rates. The importance of homeostatic plasticity may therefore parsimoniously explain both why our model fails to match the experiment, and why previous models succeeded in doing so.

The second piece is the observation that the cOSI decreases with mean firing rate in the models but not the experimental data. Importantly, selecting a threshold appropriate to the patterns of input is necessary for stimulus discriminability (Iatropoulos et al., 2024; Iatropoulos et al., 2025). In-vivo, activity-dependent mechanisms allow neurons to maintain sensory function across a range of firing rates (Brown et al., 2019), while in the models orientation selectivity is clearly rate-dependent. Not all neurons in mouse VISp respond equally robustly to parameteric grating stimuli (de Vries et al., 2020), many are optimally driven by more complex stimuli (Fu et al., 2023), and numerous variables besides stimulus modulate activity (Musall et al., 2019; Stringer et al., 2019). However, in our model and previous models neurons are specifically designed with responses to drifting gratings in mind and consequently have very similar receptive fields. Due to this, in the model, orientation selectivity is determined by how well the excitability and input of a neuron are aligned, while in-vivo it additionally reflects differences in the kinds of stimuli a neuron responds to.

Because the other variables influencing firing rate are not present in our model, and the variability of firing rates in experiments may be partly attributable to this, it is possible that the distribution of firing rates in our model is too strongly skewed, which would also be consistent with the idea that homeostatic plasticity is an important missing mechanism.

Given that previous models face this issue to a lesser degree, it may be that once it is rectified the level of orientation tuning in the models will exceed that in the experiments. Furthermore, the incorporation of other missing mechanisms could amplify it further. This is to be expected, as the models have been specifically constructed to reproduce orientation and direction tuning, with little attention to other features, while in-vivo mouse visual cortex neurons have highly complex response properties (Fu et al., 2023). This may explain why, when optimized with gradient descent, a previous visual cortex model performed unrealistically well in an orientation discrimination task (G. Chen et al., 2022). We therefore expect more complex receptive field properties, as produced by plasticity in-vivo, to be necessary to reproduce more realistc and diverse stimulus representations.

The differences between morphology-based connectivity and simpler connectivity schemes include differences in degree-distribution and topological structure. Our balancing algorithm implicitly accounted for the effect of in-degree in its conductance estimates, but nonetheless failed to generalize to the full network, indicating that topological structure, more than the indegree distribution, drives the divergence in firing rates. Consistent with this, Ito et al., 2026 reproduced strong OSI for some cell types in a network with both distance-dependent connectivity and realistic in-degree distributions (before further optimization took place).

The failure of our attempt to approximate the effects of homeostatic plasticity in section 2.9 illustrate the importance of network effects in this homeostatic balance. We propose that for realistic dynamics and stimulus encoding in detailed networks homeostatic plasticity must be simulated on-line, so that network effects can be accounted for in their operation. As we could not devise a computationally tractable way to do so, we were unable verify that this is the case. Efficiently arriving at the balanced outcome of homeostatic plasticity in detailed models is posed as a challenge for future research. The application of new computing methods, such as GPU acceleration may soon allow detailed simulation of plasticity at the necessary timescales. Alternatively, it may be more efficient to develop novel multi-scale simulation methods which would allow plasticity to be simulated with a simpler model and the results imputed onto biologically detailed one. Additionally, adaptive thresholding methods from machine learning (Y. Chen et al., 2022; Shaban et al., 2021) could be used to balance networks quickly in a non-mechanistic way. Furthermore, recently developed deep-learning based optimization methods may also make it feasible to balance complex networks while preserving biological constraints (Ito et al., 2026). Other explanations for the observed differences between our model, previous models, and experiments are possible. In appendix S4 we rule out reduced orientation or phase dependent connectivity, absence of spike-threshold adaptation, presence of short-term synaptic plasticity as explanations for the difference with previous models. We also rule out over-estimated thalamocortical innervation strength as an explanation for reduced OSI in the model. However, this does not rule out the possibility that it is some other aspect of our model which is responsible for the differences in orientation selectivity. Due to time and resource constraints, we did not validate against every possible source of data, and the validations we did performed revealed some discrepancies.

Discrepancies between our model and the experimental data which could concievably have an effect on orientation selectivity include the higher overall connection probabilities than in the MICrONs data, exaggerated distance-dependence (supplementary figure S10), innacuracies in layerwise inhibitory connection probabilities (supplementary figure S7), or low mean firing rates (while our model’s rates were plausible with respect to the data, the plausible range is large and our model is at the lower edge of it). The increased distance-dependence could conceivably limit the effect of orientation and phase dependent connectivity from distant neurons. However, if this were a major driver of low OSI in the model, we would expect phase or orientation dependent scaling to offset this and show a significant improvement in OSI, which was not the case (supplementary section S4). The remaining discrepancies would effect input-threshold balances at the population level, with some cell types receiving too much or too little excitation or inhibition. Given our results demonstrating the importance of network effects for divergence of firing rates, and previous work exploring the effects of topological complexity on network activity, we would expect that rectifying any population-level imbalances will be insufficient to produce realistic OSI without also including homeostatic plasticity.

While our results provide evidence for the importance of homeostatic plasticity in reconciling realistic connectivity with realistic response properties, it may not be the only relevant factor. Other mechanisms which could enhance orientation selectivity and therefore may be required to reproduce it in realistic networks include orientation-tuned longrange cortico-cortical feedback and direction-dependent offset in excitation and inhibition (Rossi et al., 2020).

### 3.5 Concluding remarks

Throughout this study, we attempted to reproduce characteristics of mouse VISp through detailed modeling. The failures to replicate key statistics are instructive, in that they illustrate the importance of factors not included in the model. We conclude, conditioned on the assumptions discussed, that:

1. Factors besides axo-dendritic anatomy determine the innervation of different cell types by the projection from LGd to VISp.
2. Anatomical segregation of ON/OFF inputs to the cortex plays little to no role in determining the stimulus preferences of neurons in the mouse visual cortex
3. Orientation-selective receptive field structure and like-to-like connectivity are insufficient to quantitatively explain orientation tuning in realistic networks
4. Network effects strongly influence the input-output balance of single neurons, and must be accounted for by homeostatic mechanisms

Our attribution of these discrepancies to specific missing mechanisms is speculative until those mechanisms are identified in-vivo, implemented, and demonstrated to have sufficient effect. We therefore propose the following hypotheses for further research.

1. Synaptic competition between local corticocortical, longrange corticocortical, and thalamocortical synapses mediates the innervation probability and strength to different layers in the thalamocortical projection
2. Sst and Vip expressing neurons differ from other cortical neurons in the expression of some molecular signal which determines the formation of synapses with thalamocortical afferents
3. topologically complex connectivity disrupts computation when not tempered with dynamic balancing mechanisms
4. Homeostatic plasticity is essential for orientation selectivity in the visual cortex

Realistically simulating long-term plasticity in large brain circuits is presently computationally infeasible, but the importance of plasticity to brain functioning cannot be overstated. Developing new methods to mechanistically simulate plasticity in biologically articulated models is therefore an important challenge for computational neuroscience.

We additionally emphasize the importance of validation and verisimilitude in computational modeling. The fact that mechanisms which seemed sufficient in simpler models proved insufficient when the model was made more realistic illustrates that successfully reproducing a phenomenon is not sufficient to explain it. The explanation must also be demonstrably consistent with as broad a range of evidence as possible before being relied upon. This also illustrates how it cannot be assumed that results from previous models using a given approach will hold when that approach is refined, demonstrating the need to regression test models when they are changed. The issues we faced with biophysical neuron models additionally highlight this, as calibration with respect to a broader range of features could have allowed the neuron models to generalize better to our usecase. Simulation work should carefully scrutinize as many aspects of a model as feasible, and new methods should be developed to expand this feasibility.

Additional limitations which we did not consider relevant to any particular conclusions are documented in appendix S2.

## 4 Methods

### 4.1 Cortical reconstruction

To reconstruct the anatomy of the cortex we used the methods previously described in Reimann, Bolaños-Puchet, et al., 2022, with a few key modifications.

#### 4.1.1 Circuit atlasing and composition

For the circuit geometry we used the volumetric data from the Blue Brain cell atlas (Erö et al., 2018; Piluso et al., 2025; Rodarie et al., 2021).

We enhanced it with orientation vectors generated with the regiodesics algorithm that determine the axis spanning the depth of the cortex (Gvaert, Soplata, et al., 2024). The depths of each layer along this axis were determined by intersections of rays cast along the orientation vectors with meshes representing the layer boundaries (Gvaert, Arnaudon, et al., 2024).

The ray-mesh intersections could be used to determine positions where the algorithm failed to find a realistic orientation. An orientation was considered invalid if any of the following occurred:

1. The intersecting vector approached a mesh from the wrong side
2. the vector failed to intersect with one of the layer boundaries
3. The thickness of one of the layers along the vector is more than twice that observed in mouse somatosensory cortex
4. a layer boundary is intersected more than once

These criteria marked just under 4% of voxels throughout the neocortex, largely concentrated above the corpus callosum. For these voxels both the orientation and layer boundaries were interpolated from the nearest valid neighbors.

Neuron density estimates and excitatory/inhibitory ratios in the atlas differed a greatly from experiments in the visual cortex, suggesting a limitation of its methods when applied to the visual cortex. Therefore we opted to use a different source for the cellular composition.

For the cell composition we opted instead to draw data from Schneider-Mizell et al., 2023. We used the method `scipy.stats.gaussian_kde` (Virtanen et al., 2020) to extract depthwise density profiles for each cell type in the dataset. Subsequently we mapped each of these cell types to the morpho-electrical-types (ME-types) used in our reconstruction workflow. For inhibitory cells we first mapped the cell types onto genetic markers, and then mapped those markers to ME-types. For the excitatory cells we mapped the cell types directly to ME-types.

We assumed that Sparse-targeting cells express Lysosomal Associated Membrane Protein Family Member (Lamp5), distal-targeting cells express Somatostatin (Sst), perisomatic-targeting cells express Parvalbumin (Pvalb), and Interneuron-targeting cells express Vasointestinal peptide (Vip) We then used a probabilistic mapping Roussel et al., 2021 to convert the marker-based density profiles to ME-type profiles.

For excitatory types we found that our automated classification methods performed poorly, most likely owing to errors in the skeletonization. We therefore manually established a mapping by manually inspecting exemplars from the different classes, analyzing the manually identified cell types reported in the study, and reading passages from the paper. The assigned mtypes can be seen in table 1. We only have a single electrical type for excitatory cells, only a mapping to morphological type was needed. Where a cell type corresponded to multiple m-types, the density was split in proportion to the occurrence of those m-types in the previous somatosensory cortex reconstruction Reimann, Bolaños-Puchet, et al., 2022. We did not model L6-wm neurons.

**Table 1:** Mapping used of Schneider-Mizell m-types to blue-brain m-types.

| Schneider-Mizell type | m-type |
| --- | --- |
| L2a | L2_TPC:B |
| L2b | L2_TPC:A |
| L3a, L3b | L3_TPC:A |
| L3c | L3_TPC:C |
| L4a | L4_TPC |
| L4b, L4c | L4_UPC |
| L5a, L5b | L5_TPC:C |
| L5ET | L5_TPC:A |
| L5NP | L5_TPC:B |
| L6a | L6_UPC |
| L6b | L6_TPC:C, L6_IPC |
| L6c | L6_TPC:C, L6_BPC, L6_HPC |
| L6CT | L6_TPC:A |

The resulting neuron densities followed a similar pattern to Schüz and Palm, 1989, but were somewhat higher. As the MICrONs data is a small sample from a single individual, and scaled overall density to match the numbers of Schüz and Palm, 1989, which averages 3 subjects.

For the circuit geometry and cellular composition we used the volumetric data for the primary visual cortex available from the Blue Brain cell atlas. The densities of different neuron types were used to place neurons in the CCFv3 space and assign them morpho-electrical types (METypes).

#### 4.1.2 Morphology synthesis

Instead of placing whole cellular morphologies we computationally synthesized the dendrites of neurons to match the local cortical geometry, as described in Kanari et al., 2022. Axons were assigned to these cells with the same placement rules as in previous models Markram et al., 2015; Reimann, Bolaños-Puchet, et al., 2022.

### 4.2 Thalamocortical reconstruction

The algorithm used for the thalamocortical connectivity can be summarized as follows:

1. Create straight trajectories of thalamocortical axons
2. Sample synapses on dendrites along the depth of the cortex according to a depthwise density profile. Sample more synapses than needed, for later pruning
3. Assign each synapse a presynaptic thalamocortical axon based on distance. The probability of assigning an axon to a synapse scales as a gaussian function of the distance between the synapse and the axon trajectory
4. Remove weaker connections to reproduce the original target synapse density. The probability of removal for a connection is equal to the cumulative distribution of a normal distribution evaluated at the synapse density. The variance of this distribution was fixed at 1.0 and the mean calculated for each mtype to reproduce the target fraction of pruned synapses.

For more detail, see the repository: https://github.com/BlueBrain/projectionizer/

**Table 2:** inputs of thalamocortical connectivity algorithm.

| parameter | description | experimental data |
| --- | --- | --- |
| $\sigma$ | Spread of the gaussian function used to assign axons to synapses. Represents the horizontal spread of thalamocortical axon | ballpark estimate of 120 $\mu m$ from (Reimann, Bolaños-Puchet, et al., 2022) |
| oversampling | Ratio of additional synapses to sample and subsequently prune. Increasing this will increase the number of synapses per connection and reduce connection probability and convergence. | None<br>Previously fit to synapses per connection estimates from pair recordings, now considered unreliable due to the possibility of multivesicular release (Barros-Zulaica et al., 2019) |
| cutoff variance | variance of gaussian cumulative density function to use for synapse pruning | fixed at 1.0 for sharp cutoff |
| synapse density | Number of thalamocortical synapses per unit volume at each cortical depth. | Estimated by combining (Kloc & Maffei, 2014; Meyer et al., 2010; Okigawa et al., 2021; Rodriguez-Moreno et al., 2018) (see methods) |
| axon count | total number of thalamocortical axons arriving at the visual cortex. | set at 18500 from the core of the LGN and 6000 from the shell for a total of 24500 axons. Based on excitatory neuron counts from an earlier version of the Blue Brain atlas (Piluso et al., 2025) and rounded to the nearest 100. |

#### 4.2.1 Calculation of core and shell synapse density

To calculate the depthwise profile of synapse density for the core and shell projections we combined the measurements of Kloc and Maffei, 2014 and Okigawa et al., 2021. Each of these studies provides the relative strength of fluorescence of thalamocortical axons at different depths. Kloc and Maffei, 2014 provides fluorescence for the whole of LGd, while Okigawa et al., 2021 provides density of synapses for the core only. We assume that the fluorescence scales approximately linearly with overall density of axons in these experiments (unit length per unit volume). Under this assumption we can derive the relative axon density of LGd-shell axons by subtracting the density of LGd-core axons from LGd-shell axons. We additionally assume that the peak axon density in the center of layer 4 for both studies was entirely driven by LGd-core axons, allowing us to assume that the peak density in both studies is approximately the same.

Once the values are extracted from the figures and digitized, it is clear that the peak is significantly wider in Okigawa et al., 2021. This likely reflects a difference in the specifics of how the two studies plot the fluorescence curves. To correct for this we translated the curve derived from Okigawa et al., 2021 downwards to match that derived from Kloc and Maffei, 2014.

We assume that the density of boutons scales linearly with the axon density, and that the peak bouton density is the same as in rat barrel cortex (4 × 10^7^*mm^−^*^3^ (Meyer et al., 2010)), the number of synapses per bouton is the same as in the rat POm projection (1.6 (Rodriguez-Moreno et al., 2018)), for a peak synapse density of 6.4 × 10^7^*mm^−^*^3^.

#### 4.2.2 Thalamocortical axon trajectories

The numbers of core (6000) and shell (18500) fibers were based on the total excitatory cell counts in these regions from one hemisphere of the atlas. This reflects an assumption that every excitatory neuron in the dorsal lateral geniculate nucleus sends an axon to the primary visual cortex.

Target positions for the thalamic axons in layer 4 were selected randomly from the volumetric data. At these points the principal axis (the axis pointing from white matter to pia) was retrieved from the atlas, and was used to determine the direction of the axon trajectory.

#### 4.2.3 Structural validation and adjustment

model plausibility for the fraction of cells innervated was assessed with a binomial test. The model output was assumed to accurately represent the model’s overall connection probability, allowing the model plausibility to be expressed as the p-value calculated by the method `scipy.stats.binom_test` (Virtanen et al., 2020).

model plausibility for receptive field size and relative innervation strength was assessed by a Mann-Whitney U-test (`scipy.stats.manwhitney`), where the p-value represents the probability of the sample distributions diverging at least as much as observed under the assumption that the true distributions are the same. The p-value of this test is used as the model plausability score. For the fraction of excitation per connection the model plausibility was assessed by a bootstrapping method. Measurements produced by the model were repeatedly (10000 times) sampled with a sample size (13) equal to the experiment (Lien & Scanziani, 2013). The deviation of each sample mean from the overall mean was calculated. The fraction where the deviation was greater than or equal to the experiment represents the model plausibility.

Comparing the receptive field predicted by the thalamocortical connectivity requires the assumption that this receptive field is formed by the combined receptive fields of thalamocortical afferents alone. Outside of layer 4 this assumption is unlikely to hold, as these layers receive strong local and long-range feedback (Liu et al., 2024; Shen et al., 2022; Zhang et al., 2014). Therefore, for the purposes of comparing parameter combinations we use only layer 4 pyramidal and parvalbumin-expressing cells.

#### 4.2.4 Receptive field size

To approximate the procedure of Niell and Stryker, 2008 we determined the receptive field centers of the thalamocortical afferents with a retinotopic map and partitioned input into each cell into a grid of 4 degree by 4 degree zones. We applied a gaussian blur (*σ* = 2.5*^◦^*) to the input representing the spread of the receptive fields of the thalamocortical cells and took the half-width of the region in which the resulting intensity was half-maximal or greater.

#### 4.2.5 Retinotopic map

To construct the retinotopic map of the cortex we used a flatmap of cortex (Bolaños-Puchet et al., 2024) and downsampled it to 25 micron resolution to match our atlas. We subsequently mapped the range of the visual field reported in Hübener, 2003 (from −25 to 125 degrees azimuth, from −25 to 25 degrees elevation) onto the flat space.

Receptive field centers of LGd neurons were determined by sampling the volumetric flatmap at the origin points of their axon trajectories in layer 4 (see section 4.2.2). Receptive field centers of cortical neurons were determined by sampling the volumetric flatmap at the centers of their soma.

#### 4.2.6 ON-OFF mosaic

We placed dipole centers on a flat plane using Poisson-disc sampling with a minimum distance of 24*µm* as observed in (Tring & Ringach, 2022). We then mapped this flat space onto the flatmap of the cortex (Figure 5b). We placed twice as many OFF poles as ON poles, consistent with the ratio of OFF to ON RGCs (Qiu et al., 2021), and the proportion of OFF to ON RGCs in the LGd model used Dai et al., 2020. We assigned each axon the response type of the nearest pole based on the flatmapped position of the L4 origin point of its trajectory.

#### 4.2.7 Simulating LGN input

We assigned each thalamocortical afferent a filter-based cell model matching its response class using the bmtk package (Dai et al., 2020). We used the model parameters released alongside Billeh et al., 2020. We assigned direction selective cell models to afferents originating in the shell of the LGd, and *α*-type (ON, OFF) cell models to those originating in the core of the LGd, consistent with the segregation of these pathways (Cruz-Martín et al., 2014; Okigawa et al., 2021). For direction and orientation selective cells we sampled their preferred direction from a distribution derived from Kondo and Ohki, 2016, showing the bias for cardinal directions characteristic of LGd neurons.

#### 4.2.8 Input strength calibration

To calibrate the overall strength of thalamocortical input we scaled the maximum conductance of synapses so that the mean thalamic current into L4PCs during drifting grating stimulation is approximately (46 pA) (Lien & Scanziani, 2018).

We simulated the model with drifting gratings stimulation at maximum contrast and temporal frequency of 2, with no local or background input. We recorded the total current at the soma with an in-silico voltage clamp at −70*mV*.

We then scaled all synaptic conductances by a factor equal to the experimental current divided by the current observed in the model. We ran the simulation again, and confirmed that the mean current was 46pA.

We repeated this procedure for both the biophysical and point-neuron versions of the network.

#### 4.2.9 Assessment of orientation selectivity of frequency modulation

For the experiments in section 2.3 the amplitude of modulation of the input current at the grating frequency was assessed by fitting a sine function of the stimulus temporal frequency to the signal, with offset and amplitude as free parameters. The amplitude of the resulting fit was taken as the frequency modulation amplitude. This procedure closely mimics that of Lien and Scanziani, 2013. We then calculated the orientation selectivity index of the frequency modulation amplitude (Section 4.9)

#### 4.2.10 Application of selective receptive fields

We applied the receptive field constraints calculated by Billeh et al., 2020 in our model while retaining the connectivity statistics we predicted. This was accomplished by leaving all synapses unmodified save for their source neuron. The connections were first split by whether their source node is in the core or the shell of LGd. Next, for each source and target neuron it is assessed whether the source neuron belongs in the receptive field of the target neuron. For core connections this is based on whether the source neuron’s receptive field center is located in the appropriate sub-area of the receptive field, as in Arkhipov et al., 2018. For orientation or direction selective shell neurons connections are permitted if the receptive field center is within 20 degrees of the target neuron and has a tuning angle that differs by at most 45 degrees from the target neuron.

Unfortunately this method costs some morphological specificity. Before this manipulation a synapse between a thalamocortical axon and synapse was more likely to be located closer to the axon on the dendritic tree, and therefore cells with dendrites that pass close to each other, which are therefore more likely to be connected to one another, were also more likely to receive input from the same thalamocortical axon. This relationship may play a role in reliable encoding of input signals Nolte et al., 2019. After this manipulation the relationship between synaptic location and axon location has been removed, reducing the extent to which connected neurons will share thalamic input.

#### 4.2.11 Adjusting thalamocortical short-term plasticity

In preceding cortex models (Markram et al., 2015; Reimann, Bolanõs-Puchet, et al., 2024) thalamo-cortical connections to inhibitory neurons were generally more facilitating than to excitatory neurons. We initially re-used these synaptic parameters and saw exceedingly high firing rates for layer 4 parvalbumin-expressing neurons. We therefore recalibrated the short-term dynamics of the local connectivity using data from Kloc and Maffei, 2014. We fit `tsodyks2_synapse` synapse models to relative response magnitudes in a 10Hz train of 5 spikes from Kloc and Maffei, 2014 using `scipy.optimize.differential_evolution` (Figure 14).

**Figure 14:**
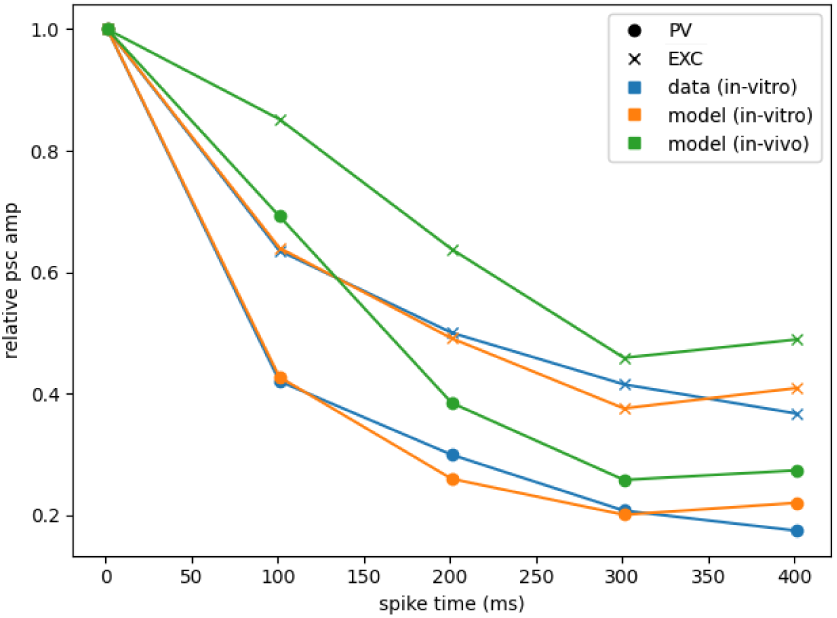
relative amplitude of responses to 10Hz spike train, for Kloc and Maffei, 2014, invitro model, and in-vivo model. The in-vitro model shows a good fit to the data, and the in-vivo model shows reduced synaptic depression as expected, while still demonstrating some depression, consistent with in-vivo studies.

Kloc and Maffei, 2014 performed their experiment in-vitro with a calcium concentration of 2mM. Initally we used Hill coefficients calculated previously (Markram et al., 2015) to adapt this to 1.3 mM as in Ecker et al., 2020, but this resulted in excessive L4PV activity and the loss of the transient activity peak at stimulus onset (compare Figure 10b and Figure S2).

We therefore instead used Jia et al., 2004 to calibrate the adjustment to in-vivo values. They measured local field potentials, which confounds the measurement of short-term synaptic plasticity by summing currents onto excitatory and pv-expressing neurons, currents from local excitatory and GABAergic synapses. We therefore fit our model to the Biculine-treated condition to avoid GABAa confounds, and fit to paired-pulse depression ratios with intervals 100ms or below to avoid GABAb confounds. This fit still does not account for possible confounds from spike-frequency adaptation or short-term dynamics of local synapses. Chung et al., 2002 suggests that at least in the barrel cortex, local synaptic depression has a minimal role in the depression of the initial stimulus response. This leaves only spike-frequency adaptation. As our model lacks spike-frequency adaptation as well, any over-estimate of synaptic depression will be compensated for by the absence of SFA.

Because this estimate comprises both synapses onto excitatory and PV-expressing neurons, we performed our calibration using a weighted sum of LGd-to-EXC and LGd-to-PV conductances. We kept the facilitation and depression time constants the same as in-vitro and varied the release probability. The three parameters for the fit were therefore the release probabilities of the two pathways and the weight of the relative contribution of LGd-to-EXC synapses. We fit them using `scipy.optimize.differential_evolution` with maximum release probabilities equal to the in-vitro case, and a minimum LGd-to-EXC weight of 0.6. Although the quality of the fit was very poor, the qualitative behavior of the synapses matched expectations, so we opted to use them nonetheless. The resulting short-term plasticity dynamics can be seen in Figure 14. The fact that a good fit could not be obtained with the fixed time constants may suggest that adjusting release probability alone is not adequate to predict in-vivo synaptic dynamics from in-vitro. As both vesicle docking and the transient increases in release probability are believed to depend on calcium concentration (Kusick et al., 2022; Zucker & Regehr, 2002), both may be affected by extracellular calcium concentration.

Importantly, in matching the strength of thalamocortical inputs to EXC and PV neurons to Ji et al., 2016, we used the product of synaptic weight and release probability. We used the in-vitro release probability, as both Kloc and Maffei, 2014 and Ji et al., 2016 used a calcium concentration of 2mM.

#### 4.2.12 Biophysical neuron models

##### Electrophysiology Data

We used the intracellular somatic whole-cell patch clamp electrophysiological recordings from P14-P16 rat somatosensory cortex excitatory and inhibitory neurons. The data were classified into 11 electrical firing types (e-types) according to the Petilla classification Ascoli et al., 2008: one excitatory (cADpyr) and 10 inhibitory: burst-accommodating (bAC), burst non-accommodating (bNAC), continuous accommodating (cAC), continuous non-accommodating (cNAC), delayed non-accommodating (dNAC), burst irregular firing (bIR), continuous irregular firing (cIR), burst stuttering (bSTUT), continuous stuttering (cSTUT) and delayed stuttering (dSTUT).

We used voltage responses from depolarising and hyperpolarising step current injections: The protocols used were IDRest, IV, APWaveForm and IDhyperpol for e-model construction. We used relative amplitude with respect to the rheobase to extract the electrical features (e-features) from protocols, e.g. IDRest_150 represents the IDRest protocol for 150% of the rheobase value. Table S1 shows the protocols, amplitudes and features extracted for these protocols. Further details about the data can be found in Reva et al., 2023 and Markram et al., 2015.

##### Morphologies

We used a scaled version of rat 3D reconstructed morphologies (Reva et al., 2023) for excitatory and inhibitory neurons. The dendrites, axons and soma of rat morphologies were scaled by a factor of 0.6622. The morphological type (m-types) of different cortical layers were used for building the e-models:

##### Inhibitory and Excitatory m-types

###### E-Models Construction

We created 5 excitatory (cADpyr) electrical models (e-models) each for cortical layers 2, 3, 4, 5 and 6: cADpyr_L2, cADpyr_L3, cADpyr_L4, cADpyr_L5, cADpyr_L6, and 10 inhibitory firing e-models corresponding to each e-type: bAC, bNAC, cAC, cNAC, dNAC, bIR, cIR, bSTUT, cSTUT and dSTUT.

We used our in-house e-model building pipeline software, BluePyEModel Mandge et al., 2025 to build the e-models which includes features extraction from patch clamp data via BluePyEfe Rössert et al., 2024 and eFEL Ranjan et al., 2024, and e-model optimisation and validation using BluePyOpt Van Geit et al., 2016.

###### Feature Extraction

The patch-clamp recordings from different were grouped into different e-types for different relative amplitudes and e-features were extracted to fit various features such as firing frequency, action potential amplitude, voltage sag, etc. A mean and standard deviation

| Inhibitory m-types | Excitatory m-types |
| --- | --- |
| DAC: Descending Axon Cell | IPC: Inverted PC |
| HAC: Horizontal Axon Cell | BPC: Bipolar PC |
| LAC: Large Axon Cell | HPC: Horizontal PC |
| NGC-DA: Neurogliaform Cell with dense axon | TPC:A: Tufted PC, late bifurcation |
| NGC: Neurogliaform Cell | TPC:B: Tufted PC, early bifurcation |
| NGC-SA: Neurogliaform Cell with sparse axon | TPC:C: Tufted PC, small tuft |
| SAC: Small Axon Cell | UPC: Untufted PC |
| BP: Bipolar Cell | SSC: Spiny Stellate Cell |
| BTC: Bitufted Cell |  |
| CHC: Chandelier Cell |  |
| DBC: Double Bouquet Cell |  |
| LBC: Large Basket Cell |  |
| MC: Martinotti Cell |  |
| NBC: Nest Basket Cell |  |
| SBC: Small Basket Cell |  |

value is calculated for each feature and amplitude using traces for the same amplitude recorded from different cells. We use a tolerance value of 10 % to select amplitudes e.g. features extracted APWavform_200 will include traces from 190% to 210 % of relative amplitude. Table S1 shows the features extracted for different amplitudes.

###### Optimisation and Validation

We built multicompartmental e-models using 3D reconstructed morphology in which the original axon was replaced by a 60 *µ*m stub axon with two axonal sections followed by a 1000 *µ*m myelinated axon. Some common parameters used for emodels are temperature = 34 *^◦^*C, specific membrane capacitance (cm) = 1 S/cm^2^, (for cADpyr: apical and basal cm = 2 S/cm^2^ to compensate for contribution for spines), initial intracellular calcium concentration = 65 * 10*^−^*^9^ M. Nernst potentials of ions: sodium (ena) = 50 mV, potassium (ek) = −90 mV. The ion channels used in model and their distributions over axonal, somatic, basal and apical dendrites as described in Reva et al., 2023.

We used the CMA-ES (Covariance matrix adaptation evolution strategy) evolutionary algorithm implemented in BluePyOpt Damart et al., 2020 to perform multiobjective optimisation. Model parameters were optimised to fit the e-features extracted from experimental data. The objective function of the optimisation was to minimise the sum of the scores of all the e-features. An e-Feature score was calculated as:

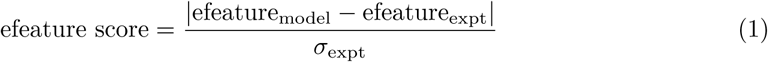

where e-feature_model_ and e-feature_expt_ are the e-feature values from optimisation and experiments, respectively, and *σ*_expt_.

The e-model was optimisation and validation involves the following steps: : i) run the protocols for resting membrane potential (RMP), input resistance (Rin), and rheobase calculation followed by other optimisation protocols, ii) calculate and minimise the features scores and iii) run the validation with the parameter set from optimisation and check if the each validation feature is below threshold score of 5. Multiple optimisations were launched with different starting seed values for the optimiser. The optimisation with the lowest value which was validated was selected.

**Table 3:** Parameterization of point models.

|  |  |
| --- | --- |
| parameter | provenance |
| threshold potential | set to reproduce the rheobase from the experiment,<br>$V_{th} = V_l \frac{I_{th}}{g_l}$ |
| resting potential | directly from database |
| leak conductance | inverse of input resistance |
| capacitance | leak conductance divided by the membrane time constant |
| reset potential | assumed to be the same as resting potential |

### 4.3 Conversion to NEST

In order to reproduce orientation selectivity we converted our model to the NEST simulator (Gewaltig & Diesmann, 2007). For each neuron in the model we selected a neuron from the Allen Institute Cell Types Database (Gouwens et al., 2019). We used the model neuron’s assigned layer and mclass to identify appropriate experiments to sample from. As the database does not have measurements of Lamp5-expressing neurons, we used experiments with neurons expressing 5Htr3a for this m-class.

We used the `glif_cond` neuron model type. We did not make use of the threshold-adaptation capabilities of the GLIF neuron models, as these were not necessary for our research questions and would have limited our sample size of model parameters to neurons for which models had been successfully fit, thus also potentially introducing new sampling biases.

All connections were reduced to single `tsodyks2_synapse synapse models`. In multisynapse models all synaptic parameters were averaged across synapses for the condensed connection. Synaptic delay was averaged across synapses in the connection. It should be noted that while this preserves delay due to axonal propagation, delays due to dendritic integration are lost using this approach.

After converting the model to NEST-compatible form we scaled the synapses so that the first-spike synaptic conductances for each pathway matched those in the Allen Institute Synaptic Physiology dataset Campagnola et al., 2022. To account for distance-dependent effects, for this calibration we binned the experimental data by the horizontal distances between neurons in bins of 25*µm* and sampled model connections from each distance bin in proportion to the number of samples in the experimental dataset. We additionally scaled synaptic release probabilities for all pathways to account for the differences in calcium concentration (1.3mM in Campagnola et al., 2022 and 2.0mM in the experiments used to calibrate the connectivity previously) as in Markram et al., 2015. Differences in release probability can affect the post-synaptic current amplitudes recorded in pair recordings. We therefore adjusted synaptic weights in inverse proportion to release probability 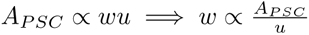.

Initial simulations for firing rate calibration showed that most neurons were silent even when mean firing rates were high, while a few neurons had extremely high firing rates. As a first effort at addressing this, we ranked the neurons within each m-class by total afferent synaptic weight and reassigned electrical models to them according to their rank, such that the electrical models with the highest rheobase for a given m-class are assigned to the neurons with the greatest synaptic input. This mostly rectified the issue, and the resulting distributions are plausible. Simulation results shown are from models with sorted electrophysiology.

### 4.4 Local connectivity recipe

In previous work, we underestimated inhibitory connection probability, a risk noted in the original paper (Reimann et al., 2015) due to using synapses-per-connection estimates from pair recordings rather than electron microscopy data, which would over-estimate synapse counts in the presence of multi-vesicular release. We updated the algorithm with new synapses-per-connection data from Schneider-Mizell et al., 2023. Aside from this, the original synaptic connectivity recipe was left unchanged from the rat somatosensory cortex model of Reimann, Bolanõs-Puchet, et al., 2024.

### 4.5 Rewiring connectivity

#### 4.5.1 Calculating conditional synapse count distribution from MICrONS

We re-calibrated the conditional synapse count distribution using a pre-processed version of the MICrONs dataset, available at https://doi.org/10.5281/zenodo.13849415.

From the MICrONS dataset we have access to the distance and number of synapses (including zeros) between cells. To avoid edge-effects we restricted our analysis to a column 150 *µm* in radius. Binning the horizontal distances between neurons at 25*µm* provided us with us a matrix *S* where *S_ij_*= *P* (*n_syn_* = *j*|25*i < d <* 25(*i* + 1).

From our model we have access to the distance and number of appositions between cells. We once again restricted our analysis to a column of 150*µm* and binned horizontal distances at 25 *µm*, providing us with a matrix *N* where *N_ik_* = *P* (*n_app_* = *k*|25*i < d <* 25(*i* + 1)).

To match the different cell types, we used the Schneider-Mizell classification (Schneider-Mizell et al., 2023) of the neurons in the MICrONs dataset and mapped them onto genetic classes in the model, mapping STCs to Lamp5 neurons, ITCs to Vip neurons, DTCs to Sst neurons, PTCs to Pvalb neurons, and all remaining types to the excitatory neurons.

We assumed that the relationship between distance and connectivity is mediated by the number of appositions, such that

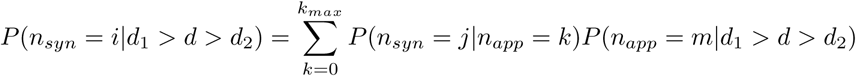

This can be alternatively written as *S* = *NA* where *A_kj_* = *P* (*n_syn_* = *j*|*n_app_* = *k*). We therefore needed to find a solution *Â* that satisfies the constraints 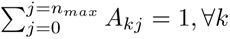, representing the fact that the rows of A represent conditional probabilities, and *j > k* =⇒ *A_jk_* = 0, representing the fact that the number of synapses must be fewer than the number of appositions.

This problem is underconstrained. To reduce the number of dimensions, we assumed that the number of synapses between a pair of neurons follows a binomial distribution with the number of trials equal to the number of appositions, and the probability of success additionally depending on the number of appositions. The formula for the probability of success is *p* = *a* ∗ *n_app_* + *b*, capped between 0 and 1.

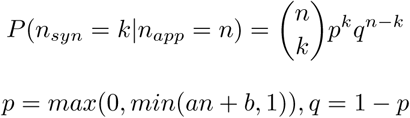

We truncated the binomial distribution at the maximum number of synapses observed in any connection. The parameter *b* represents a general synapse probability, while *a* can be tuned to adjust how strongly the connectivity depends on the number of appositions.

The sample we use to estimate *P* (*n_app_* = *k*|*d_h_*= *d*) contains only a few pairs with large numbers of appositions. Consequently, many apposition counts represented in the model as a whole do not show up in the sample. We fit an inverse binomial function to the apposition distribution at each distance *P* (*n_app_* = *n*|*d*_1_ *< d < d*_2_) and used this to determine the missing values. While this seemed to underestimate the probability of larger apposition counts, it was nonetheless sufficient for our purposes.

We optimized for a and b using the differential evolution algorithm from scipy (Virtanen et al., 2020) with the rand1bin strategy and boundaries of [−0.1, 0.1] for a and [0, 0.2] for b. The loss function was the mean square error, *L*(*Â*) = ||*N̂Â* − *Ŝ*||^2^. The resulting parameters and log-likelihood by pathway can be found in table 4.

**Table 4:**
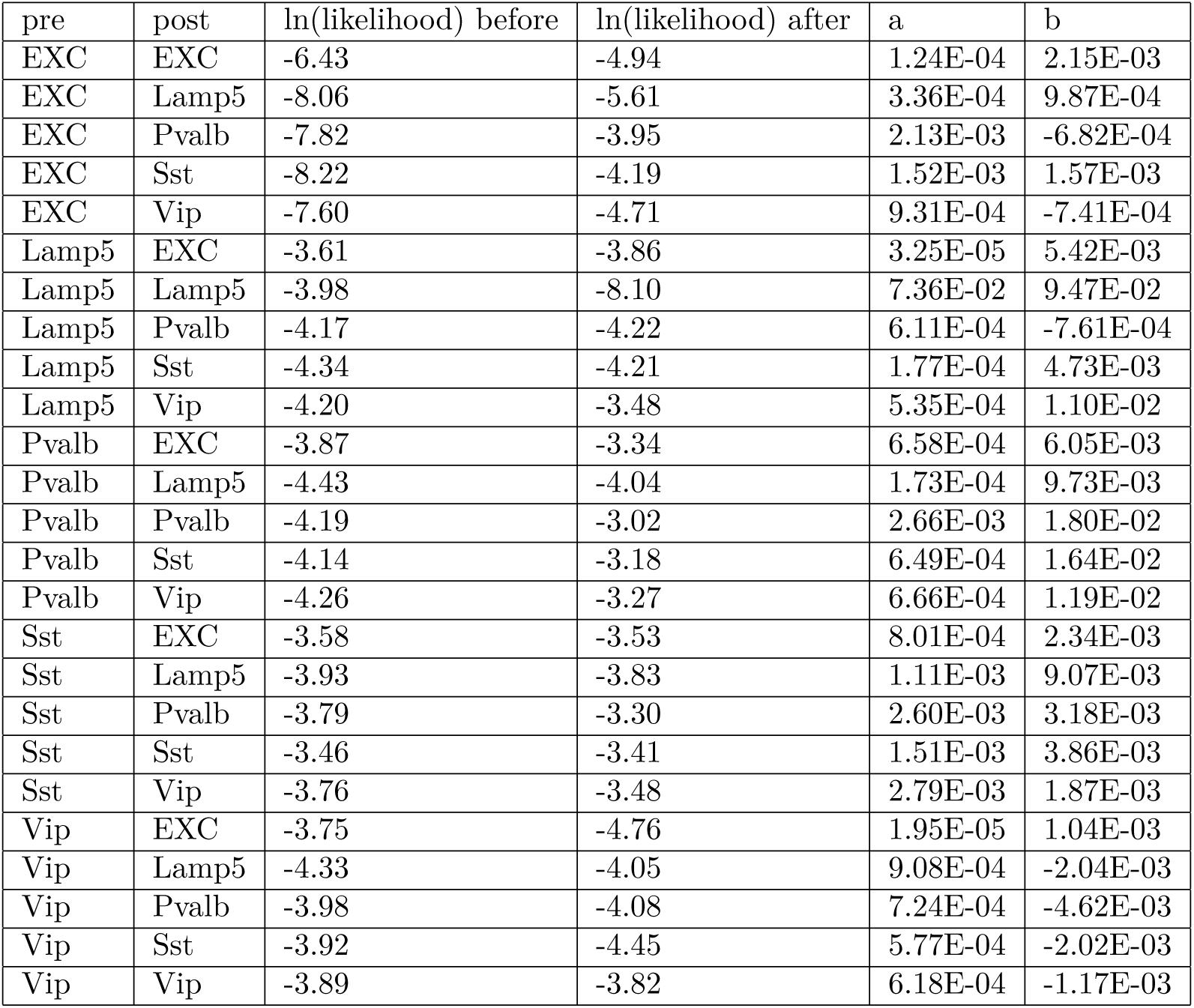
parameters and log-likelihood scores of the connectivity fit. Note that for most pathways the log-likelihood improved, with the exception of a few Lamp5 or Vip pathways, where the smaller sample size may have led to overfitting on the train set.

| pre | post | ln(likelihood) before | ln(likelihood) after | a | b |
| --- | --- | --- | --- | --- | --- |
| EXC | EXC | -6.43 | -4.94 | 1.24E-04 | 2.15E-03 |
| EXC | Lamp5 | -8.06 | -5.61 | 3.36E-04 | 9.87E-04 |
| EXC | Pvalb | -7.82 | -3.95 | 2.13E-03 | -6.82E-04 |
| EXC | Sst | -8.22 | -4.19 | 1.52E-03 | 1.57E-03 |
| EXC | Vip | -7.60 | -4.71 | 9.31E-04 | -7.41E-04 |
| Lamp5 | EXC | -3.61 | -3.86 | 3.25E-05 | 5.42E-03 |
| Lamp5 | Lamp5 | -3.98 | -8.10 | 7.36E-02 | 9.47E-02 |
| Lamp5 | Pvalb | -4.17 | -4.22 | 6.11E-04 | -7.61E-04 |
| Lamp5 | Sst | -4.34 | -4.21 | 1.77E-04 | 4.73E-03 |
| Lamp5 | Vip | -4.20 | -3.48 | 5.35E-04 | 1.10E-02 |
| Pvalb | EXC | -3.87 | -3.34 | 6.58E-04 | 6.05E-03 |
| Pvalb | Lamp5 | -4.43 | -4.04 | 1.73E-04 | 9.73E-03 |
| Pvalb | Pvalb | -4.19 | -3.02 | 2.66E-03 | 1.80E-02 |
| Pvalb | Sst | -4.14 | -3.18 | 6.49E-04 | 1.64E-02 |
| Pvalb | Vip | -4.26 | -3.27 | 6.66E-04 | 1.19E-02 |
| Sst | EXC | -3.58 | -3.53 | 8.01E-04 | 2.34E-03 |
| Sst | Lamp5 | -3.93 | -3.83 | 1.11E-03 | 9.07E-03 |
| Sst | Pvalb | -3.79 | -3.30 | 2.60E-03 | 3.18E-03 |
| Sst | Sst | -3.46 | -3.41 | 1.51E-03 | 3.86E-03 |
| Sst | Vip | -3.76 | -3.48 | 2.79E-03 | 1.87E-03 |
| Vip | EXC | -3.75 | -4.76 | 1.95E-05 | 1.04E-03 |
| Vip | Lamp5 | -4.33 | -4.05 | 9.08E-04 | -2.04E-03 |
| Vip | Pvalb | -3.98 | -4.08 | 7.24E-04 | -4.62E-03 |
| Vip | Sst | -3.92 | -4.45 | 5.77E-04 | -2.02E-03 |
| Vip | Vip | -3.89 | -3.82 | 6.18E-04 | -1.17E-03 |

We performed this calculation with a 2/3 subset of the MICrONs dataset and assessed the likelihood of the resulting conditional distributions *P* (*n_syn_* = *n*|*d_h_* = *d*) on a 1/3 subset. Overall, overfitting was minimal and the likelihood of the adjusted model was better than that of the original.

After evaluating the quality of the fit, we re-ran the calibration using the whole dataset rather than only the training subset in order to get the best possible result.

#### 4.5.2 Receptive fields for rewiring

We created a grid representation of each neuron’s thalamocortical receptive fields as described in section 4.2.4. For neurons without thalamocortical input, we used all candidate inputs that matched its assigned receptive field properties (ON/OFF, transient/sustained subfields).

#### 4.5.3 Correlation-based rewiring

To reproduce the relationships observed between connectivity and pairwise response correlation we re-assigned synapses within the model such that pairs of neurons with high response correlation are more likely to connect and preferentially received stronger connections. We conserved the effects of the axo-dendritic touches on the connectivity, and assumed all variation in connection strength unexplained by axo-dendritic touches is instead explained by response correlation. It is implemented within the connectome-manipulator framework (Pokorny et al., 2024), but unlike previous manipulations accounts for the axo-dendritic touches in the model.

First, the connectome is split into pathways between different cell types. We chose to split by layer and marker, with layers L1, L23, L4, L5 and L6, and markers Lamp5, Vip, Pvalb, Sst, and one EXC class for all excitatory neurons.

Next pairs of neurons are grouped by the number of axo-dendritic touches (as calculated by the touchdetector algorithm (Markram et al., 2015; Reimann, Bolaños-Puchet, et al., 2022)) Pairs of neurons with no axo-dendritic touches are disregarded.

For each pathway we use a matrix of conditional probabilities *A* such that *A_kj_* = *P* (*n_syn_* = *j*|*n_app_*= *k*), where *n_syn_* and *n_app_* are the number of synapses and the number of axo-dendritic touches, respectively. This matrix includes values for *n_syn_* = 0, thereby accounting for connection probability. Initially these matrices were sampled from the model in order to preserve its anatomy-connectivity relationships. Afterward these were calculated using experimental data, as explained in the preceding subsection 4.5.1.

All pairs with potential connections have their response correlation evaluated. For each postsynaptic neuron, the response correlation with potential presynaptic partners (*n_app_ >*= 1) is evaluated, ranked, and has a quantile assigned *q_r_*. This is such that the most correlated partner will have a *q_r_* value of 100%, and the least correlated 1*/n_candidates_*. This ranking is done independently for all afferents to each neuron rather than among all pairs of neurons so that total synapse counts on a neuron will be independent of its receptive field properties.

Each presynaptic partner is assigned a number of synapses 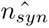. This is selected as the largest value which satisfies 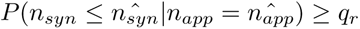. In this way all variation in the number of synapses not determined by the number of appositions is determined by the rank of the response correlation, such that the weakest connections are assigned to the least correlated pairs. As all variability is ‘explained away’ by *q_r_*, this algorithm is deterministic with respect to its inputs. Note that when the number of candidate afferents is small the smallest *q_r_* will not be very small, and this may consequently over-assign synapses, which may explain the increased connection probability seen in the model compared to the conditional distribution (Supplementary figure S10).

Subsequently all synapses within a pathway are sorted by their conductance and assigned to the sorted connections, such that synapses with higher conductance deterministically go to the pairs with higher response correlation.

Finally, synapse positions are assigned from the pool of potential synapse sites (axo-dendritic touches) between the pre and post-synaptic pairs. Synaptic delays are calculated based on the path distance along the axon to the synapse site, assuming a propagation velocity of 300*µm/ms*, as in previous models (Markram et al., 2015; Reimann, Bolaños-Puchet, et al., 2022).

#### 4.5.4 Assessment of connectivity-correlation relationship

We selected all layer 2/3 excitatory neurons within 125 *µm* of the region’s center. For all pairs we calculated the Pearson correlation of their receptive fields.

We used a bootstrapping method to assess the hypothesis that the experimental result could have been sampled from the same distribution as the model. We sub-sampled the pairs of neurons 520 at a time (sample size of experiment), calculated the proportion of synaptic strength explained by the top n% most correlated pairs (where n was either 7 or 12, depending on the comparison), and repeated this 10000 times. If the model’s predicted value was 50% or less, then the p-value was the fraction of samples greater or equal to 50% of synaptic strength. Otherwise, the p-value was the fraction of samples less than or equal to 50% synaptic strength.

For orientation-dependent connection probability, we used a binomial test to compare model connection probabilities for each bin of orientation preference difference.

### 4.6 NEST simulations

Simulations were run for neurons within 700 *µm* of the center of VISp with NEST version 3.8.0-post0.dev0 compiled with gcc 13.2.0, cmake 3.27.9, boost 1.85.0 on a single node (Intel(R) Xeon(R) Platinum 8360Y CPU @ 2.40 GHz).

Drifting grating simulations for the full model were additionally re-run with NEST 3.9.0-post0.dev0 to verify that a bug we encountered (https://github.com/nest/nest-simulator/issues/3634}) in 3.8, but not 3.9, was not responsible for the reduced orientation selectivity.

### 4.7 Circuit activity calibration

Our model does not explicitly contain long-range cortico-cortical synapses. To compensate for this we used two slightly different approaches for the biophysical and point-neuron networks.

In the biophysical network we used an Ornstein-Uhlenbeck process with a coefficient of variation of 0.4 and a mean strength scaled to the input resistance of the neuron, as in Isbister et al., 2025. In the point-neuron version, we placed one additional synapse on each neuron, connected to a presynaptic Poisson process firing at 1KHz as in Billeh et al., 2020. The synapse model `static_synapse` from NEST allows overlapping synaptic events, thereby simulating an input pattern as though from multiple synapses. The strength of this synapse was scaled in proportion to the neuron’s threshold conductance 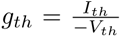 where *V_th_* is the threshold potential and *I_th_* is the rheobase. The formula for longrange synapse strength is therefore 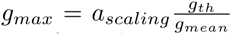 where *a_scaling_* is a cell-type specific constant and *g_mean_* is the mean conductance from the synapse at a weight of 1 and rate of 1000Hz. In effect, for *a_scaling_* = 1, the mean conductance from the synapse throughout a simulation is equal to the threshold conductance *g_th_*.

We calibrated the strength of these additional inputs to each cell class by aiming to match the firing rates observed in Siegle et al., 2021.

Siegle et al., 2021 uses extracellular electrodes to record neural activity. These methods can lead to multiple neurons being detected as a single unit (Buzsáki, 2004; “ecephys_quality_metrics”, n.d.; Wohrer et al., 2013). Such errors can be screened out by identifying and excluding units with spikes that occur unrealistically close together (ISI violations). To maximize the accuracy of the firing rate estimates we used only units where no ISI violations were detected.

Alongside this, extracellular electrophysiology also preferentially samples neurons with higher firing rates. To compensate for this we compared the firing rates observed for fast-spiking neurons in Siegle et al., 2021 and parvalbumin-expressing neurons in W.-p. Ma et al., 2010, The estimated adjustment ratio of 32% is similar to previous estimates (Laquitaine et al., 2024; Olshausen & Field, 2005).

We therefore adjusted the target firing rates by this ratio. It was necessary to assume that the same ratio applies to all cell types and layers. It should be noted however that the ratio between W.-p. Ma et al., 2010 and Siegle et al., 2021 firing rates varies between layer 4 and layer 23, indicating that this assumption does not entirely hold. Understanding the extent of sampling biases in different neural populations through simulation of extracellular recordings (Laquitaine et al., 2024; Tharayil et al., 2025) may allow for a more accurate calibration of network models. Adapting the methods of Isbister et al., 2024, we adopted a two-step optimization approach.

In the first step we simulated the model without any local connectivity and varied *a_scaling_*for all neurons between 0 and 1. Based on this we estimated the relative strength of missing excitation to each cell group *a_relative_*as the value of *a_scaling_*where the mean firing rate for that cell group is the closest to the experimental mean.

Next we simulated a central column 500*µm* across and measured firing rates only from a central column of 300*µm*. We varied the overall amount of background input such that *a_scaling_* = *a_overall_* ∗ *a_relative_*, varying *a_overall_*between 0 and 1. We compared the mean firing rates of the experiment and model at each value, and selected the value for which the mean squared error was the smallest. The experimental data did not contain any firing rates for layer 1, we therefore assumed the same *a_relative_*as in layer 2/3.

Experiments have also shown that different types of neurons and pyramidal cells receive longrange input in different amounts and from different sources (Shen et al., 2022; Vélez-Fort et al., 2014; Zhang et al., 2014). Our calibration only distinguishes betwen fast-spiking (parvalbumin-expressing) and regular spiking neurons, which is likely a source of additional inaccuracy.

### 4.8 Altered models

In appendix S4 we show orientation selectivity of slightly altered models to show that these alterations do not affect orientation selectivity.

#### 4.8.1 Orientation-dependent scaling

We quantified the diference in orientation preferences of pre and post synaptic pairs (between 0 and 90 degrees). Connection variabilty was very high in our model, so our scaling was designed to reduce variance in connection strength. We defined a parameter *p_w_*, and set synaptic weights s.t. 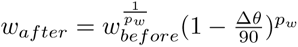. We varied *p_w_* and visualized the relationship between the difference in preference and connection strength, choosing a value of 1.3 because it exaggerated the relationship of the experiment without eliminating iso-orientation connectivity entirely. Weights were subsequently scaled to preserve the mean strength.

#### 4.8.2 Phase-dependent scaling

We scaled the connection strength linarly with phase. Phase difference Δ*p* defined as retinotopic distance projected in the neuron’s preferred direction. Phase difference was capped at 4 degrees, and weights scaled s.t. 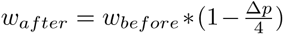. Weights were subsequently scaled to preserve the mean strength.

#### 4.8.3 GLIF neuron models

We replaced all neuron models with corresponding neuron models sampled from the model of Billeh et al., 2020.

#### 4.8.4 No short-term plasticity

We replaced all synapse models with static synapses, with weight equal to the product of the previous weight and release probability (preserving PSC amplitude). Firing rates remained similar.

### 4.9 Orientation selectivity index

The orientation selectivity index used was the global orientation selectivity index, as used in Billeh et al., 2020; Lien and Scanziani, 2013; Siegle et al., 2021. It takes the form:

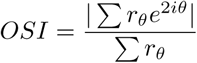

Where *r_θ_* is the response at grating drift direction *θ*. For calculating the OSI of frequency modulation we used the modulation amplitude directly. When evaluating OSI (but not cOSI, see below) of firing rates we first averaged firing rates across trials before subtracting the lowest average firing rate from all firing responses.

The control-corrected OSI (cOSI) is calculating by subtracting the OSI of a control with shuffled responses from the OSI of the model neuron. In this case, we do not subtract a baseline activity level, allowing this metric to serve as an indicator of the efficacy orientation-tuning in cell-to-cell signaling.

### 4.10 Per-neuron input calibration

We aimed to satisfy the equation:

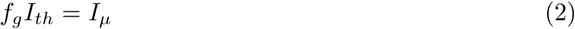

Where *I_th_* is a neuron’s rheobase, *f_g_* is a cell-type dependent coefficient, and *I_µ_* is the expected mean current at threshold. i.e.

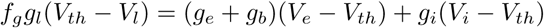

Where *g_l_, g_e_, g_b_, g_i_* are leak conductance and mean excitatory, background and inhibitory conductance, *V_th_, V_l_, V_e_, V_i_* are threshold potential, resting potential, and excitatory and inhibitory reversal potentials.

To be able to determine *g_e_* and *g_i_* for each neuron from its connectivity we ran simulations with 100 neurons of the given type, in which all afferent synapses fire with a Poisson process at rates sampled from the target distribution of firing rates for the presynaptic cell type. i.e. a connection coming from a layer 2/3 excitatory neuron will have a rate sampled from the desired firing rate distribution for layer 2/3 excitatory neurons. In effect, this simulates inputs to the simulated neurons from a perfectly calibrated network.

We recorded *g_e_* and *g_i_* for these neurons. We then summed the synaptic weights from each mclass to these neurons in matrices *W_e_* and *W_i_* for excitatory and inhibitory weights respectively. We then fit matrices *A_e_* and *A_i_* with linear regression such that *W_e_A_e_* ≈ *g_e_*, *W_i_A_i_* ≈ *g_i_*. We used cross-validation to assess the accuracy of this fit. We varied the sample size used for fitting from 10 to 80 neurons, and assessed the quality of the fit on the remaining neurons from the original sample of 100. At each sample size we repeated this 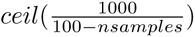 times with re-randomized train-test assignment to ensure that when the number of neurons in the test set is small, the number of reptitions are scaled up in proportion. The Pearson correlation of the predicted and actual mean conductance went above 0.99 for most neuron types, and was above 0.9 for all of them, indicating a good fit.

Having a means to predict the excitatory and inhibitory conductances from local and thalamocortical synapses, we had a means to estimate a magnitude for the mean background conductance *g_b_* to each neuron for a given value of *f_g_*.

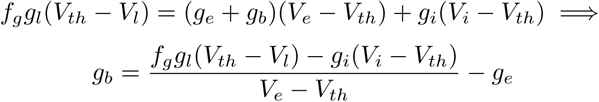

For some neurons, at some values of *f_g_*, *g_b_* would be negative. For these neurons we set *g_b_* to 0 and adjusted their threshold potential as necessary to satisfy the equation, i.e.

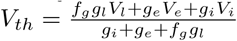

By adjusting the input conductance and threshold potentials in this way we could ensure the equation is satisfied for any value of *f_g_*. We then sampled 100 new neurons, predicted *g_e_, g_i_* and varied *f_g_*in idealized simulations. We measured their firing rates and compared them to the target firing rates, performing a bisecting search over values of *f_g_*until the difference in mean firing rates was less than 0.3*Hz*.

We then created a version of the network in which the background input to each neuron was scaled to satisfy equation 2 for the input conductances predicted based on its connectivity.

## 5 Data and code availability

Model and analysis code will be made available, at the latest, with publication of the article.

## 6 Author Contributions

Alexis and Darshan optimized the biophysical neuron models and helped in identifying the issues with them. Joni provided technical support with the thalamocortical connectivity algorithm, as well as code review. Armando and Henry provided advice on the model and feedback on the manuscript. The rest was done by Hugo.

