## Supplementary materials for "Do we understand orientation selectivity? A simulation study"

### Tables

| E-type | Protocol | E-features |
|---|---|---|
| cADpyr (L6, L5, L4, L3, L2) | APWaveform 320 % | AP_amplitude, AP1_amp, AP2_amp, AP_duration_half_width, AHP_depth |
| | IV -100 % | voltage_deflection, voltage_deflection_begin |
| | IDrest & IDthresh (Step) 130, 150, 200, 280 % | voltage_base, voltage_after_stim, AP_amplitude, APlast_amp, AHP_depth, inv_time_to_first_spike, time_to_last_spike, inv_first_ISI, inv_second_ISI, inv_third_ISI, inv_fourth_ISI, inv_fifth_ISI, mean_frequency, strict_burst_number, ISI_CV |
| | IDhyperpol 150 % | voltage_base, mean_frequency, Spikecount |
| | IV -40 % (Rin) | ohmic_input_resistance_vb_ssse, voltage_base |
| | IV 0 % (RMP) | voltage_base, Spikecount |
| | SearchHoldingCurrent | bpo_holding_current |
| | SearchThresholdCurrent | bpo_threshold_current |
| | bAP | Spikecount, maximum_voltage_from_voltagebase, maximum_ca_prox_apic_from_voltagebase, maximum_ca_prox_basal_from_voltagebase, maximum_ca_prox_soma_from_voltagebase, maximum_ca_prox_ais_from_voltagebase |
| bAC, cACint | IDThresh/IDrest (Step) 130, 140, 200, 250, 300 % | voltage_base, voltage_after_stim, AP_amplitude, APlast_amp, AHP_depth, inv_time_to_first_spike, time_to_last_spike, inv_first_ISI, inv_second_ISI, inv_third_ISI, inv_fourth_ISI, inv_fifth_ISI, mean_frequency, strict_burst_number, ISI_CV |
| | APWaveform 360 % | AP_amplitude, AP1_amp, AP_duration_half_width, AHP_depth |
| | IV -100 % | voltage_deflection, voltage_deflection_begin |
| | IV -40 % (Rin) | ohmic_input_resistance_vb_ssse, voltage_base |
| | IV 0 % (RMP) | voltage_base, Spikecount |
| | IDhyperpol 150 % | voltage_base, mean_frequency, Spikecount |
| | SearchHoldingCurrent | bpo_holding_current |
| | SearchThresholdCurrent | bpo_threshold_current |
| bNAC, cNAC | IDThresh/IDrest (Step) 130, 150, 200, 250, 300 % | voltage_base, voltage_after_stim, AP_amplitude, APlast_amp, AHP_depth, inv_time_to_first_spike, time_to_last_spike, inv_first_ISI, inv_second_ISI, inv_third_ISI, inv_fourth_ISI, inv_fifth_ISI, mean_frequency, strict_burst_number, ISI_CV |
| | APWaveform 360 % | AP_amplitude, AP1_amp, AP_duration_half_width, AHP_depth |
| | IDhyperpol 150 % | voltage_base, mean_frequency, Spikecount |
| | IV -100 % | voltage_deflection, voltage_deflection_begin |
| | IV -40 % (Rin) | ohmic_input_resistance_vb_ssse, voltage_base |
| | IV 0 % (RMP) | voltage_base, Spikecount |
| | SearchHoldingCurrent | bpo_holding_current |
| | SearchThresholdCurrent | bpo_threshold_current |
| dNAC | IDThresh/IDrest (Step) 120, 130, 150, 200, 260 % | voltage_base, voltage_after_stim, AP_amplitude, APlast_amp, AHP_depth, inv_time_to_first_spike, time_to_last_spike, inv_first_ISI, inv_second_ISI, inv_third_ISI, inv_fourth_ISI, inv_fifth_ISI, mean_frequency, strict_burst_number, ISI_CV |
| | APWaveform 300 % | AP_amplitude, AP_duration_half_width |
| | IDhyperpol 150 % | voltage_base, mean_frequency, Spikecount |
| | IV -100 % | voltage_deflection, voltage_deflection_begin |
| | IV -40 % (Rin) | ohmic_input_resistance_vb_ssse, voltage_base |
| | IV 0 % (RMP) | voltage_base, Spikecount |
| | SearchHoldingCurrent | bpo_holding_current |
| | SearchThresholdCurrent | bpo_threshold_current |

| | | |
|---|---|---|
| cSTUT | IDThresh/IDrest (Step)<br>130, 180, 280 % | voltage_base, voltage_after_stim, AP_amplitude,<br>APlast_amp, AHP_depth, inv_time_to_first_spike,<br>time_to_last_spike, inv_first_ISI, inv_second_ISI,<br>inv_third_ISI, inv_fourth_ISI, inv_fifth_ISI, inv_last_ISI<br>mean_frequency, strict_burst_number, ISI_CV |
| | IDThresh/IDrest (Step)<br>140, 240 % | mean_frequency,inv_time_to_first_spike,time_to_last_spike,<br>inv_first_ISI, inv_second_ISI, inv_third_ISI, inv_fourth_ISI,<br>inv_fifth_ISI, inv_last_ISI, strict_burst_number, ISI_CV |
| | IDhyperpol 150 % | voltage_base, mean_frequency, Spikecount |
| | APWaveform 320 % | AP_amplitude, AP1_amp,<br>AP_duration_half_width, AHP_depth |
| | IV -100 % | voltage_deflection, voltage_deflection_begin |
| | IV -40 % (Rin)<br>IV 0 % (RMP) | ohmic_input_resistance_vb_ssse, voltage_base<br>voltage_base, Spikecount |
| | SearchHoldingCurrent | bpo_holding_current |
| | SearchThresholdCurrent | bpo_threshold_current |
| bSTUT | IDThresh/IDrest (Step)<br>130, 140, 280 % | voltage_base, voltage_after_stim, AP_amplitude,<br>APlast_amp, AHP_depth, inv_time_to_first_spike,<br>time_to_last_spike, inv_first_ISI, inv_second_ISI,<br>inv_third_ISI, inv_fourth_ISI, inv_fifth_ISI, inv_last_ISI,<br>mean_frequency, strict_burst_number, ISI_CV |
| | IDThresh/IDrest (Step)<br>180, 220 % | mean_frequency, inv_time_to_first_spike,time_to_last_spike,<br>inv_first_ISI, inv_second_ISI, inv_third_ISI, inv_fourth_ISI,<br>inv_fifth_ISI, inv_last_ISI, strict_burst_number, ISI_CV |
| | IDhyperpol 150 % | voltage_base, mean_frequency, Spikecount |
| | APWaveform 300 % | AP_amplitude, AP1_amp,<br>AP_duration_half_width, AHP_depth |
| | IV -100 % | voltage_deflection, voltage_deflection_begin |
| | IV -40 % (Rin) | ohmic_input_resistance_vb_ssse, voltage_base |
| | IV 0 % (RMP) | voltage_base, Spikecount |
| | SearchHoldingCurrent | bpo_holding_current |
| | SearchThresholdCurrent | bpo_threshold_current |
| dSTUT | IDThresh/IDrest (Step)<br>120, 130, 200 % | voltage_base, voltage_after_stim, AP_amplitude,<br>APlast_amp, AHP_depth, inv_time_to_first_spike,<br>time_to_last_spike, inv_first_ISI, inv_second_ISI,<br>inv_third_ISI, inv_fourth_ISI, inv_fifth_ISI, inv_last_ISI,<br>mean_frequency, strict_burst_number, ISI_CV |
| | IDThresh/IDrest (Step) 140 % | mean_frequency, inv_time_to_first_spike, time_to_last_spike,<br>inv_first_ISI, inv_second_ISI, inv_third_ISI, inv_fourth_ISI,<br>inv_fifth_ISI, inv_last_ISI, burst_number, ISI_CV |
| | APWaveform 300 % | AP_amplitude, AP1_amp,<br>AP_duration_half_width, AHP_depth |
| | IDhyperpol 150 % | voltage_base, mean_frequency, Spikecount |
| | IV -100 % | voltage_deflection, voltage_deflection_begin |
| | IV -40 % (Rin) | ohmic_input_resistance_vb_ssse, voltage_base |
| | IV 0 % (RMP) | voltage_base, Spikecount |
| | SearchHoldingCurrent | bpo_holding_current |
| | SearchThresholdCurrent | bpo_threshold_current |
| cIR | IDThresh/IDrest (Step)<br>130, 220, 280 % | voltage_base, voltage_after_stim, AP_amplitude,<br>APlast_amp, AHP_depth, inv_time_to_first_spike,<br>time_to_last_spike, inv_first_ISI, inv_second_ISI,<br>inv_third_ISI, inv_fourth_ISI, inv_fifth_ISI, inv_last_ISI<br>mean_frequency, strict_burst_number, ISI_CV |
| | IDThresh/IDrest (Step)<br>140, 180 % | mean_frequency ,inv_time_to_first_spike, time_to_last_spike,<br>inv_first_ISI, inv_second_ISI, inv_third_ISI, inv_fourth_ISI<br>inv_fifth_ISI, inv_last_ISI, strict_burst_number, ISI_CV |
| | APWaveform 280 % | AP_amplitude, AP1_amp, |
| | IDhyperpol 150 % | voltage_base, mean_frequency, Spikecount |
| | | AP_duration_half_width, AHP_depth |
| | IV -100 % | voltage_deflection, voltage_deflection_begin |
| | IV -40 % (Rin) | ohmic_input_resistance_vb_ssse, voltage_base |

| | | |
|---|---|---|
| | IV 0 % (RMP) | voltage_base, Spikecount |
| | SearchHoldingCurrent | bpo_holding_current |
| | SearchThresholdCurrent | bpo_threshold_current |
| bIR | IDThresh/IDrest (Step) 180, 240, 280 % | voltage_base, voltage_after_stim, AP_amplitude, APlast_amp, inv_time_to_first_spike,time_to_last_spike, inv_first_ISI, inv_second_ISI, inv_third_ISI, inv_fourth_ISI, inv_fifth_ISI, inv_last_ISI, AHP_depth_abs mean_frequency |
| | IDThresh/IDrest (Step) 140, 200 % | mean_frequency, inv_time_to_first_spike,time_to_last_spike, inv_first_ISI, inv_second_ISI, inv_third_ISI, inv_fourth_ISI inv_fifth_ISI, inv_last_ISI, burst_number, ISI_CV |
| | APWaveform 280 % | AP_amplitude, AP1_amp, AP_duration_half_width, AHP_depth_abs |
| | IDhyperpol 150 % | voltage_base, mean_frequency, Spikecount |
| | IV -100 % | voltage_deflection, voltage_deflection_begin |
| | IV -40 % (Rin) | ohmic_input_resistance_vb_ssse, voltage_base |
| | IV 0 % (RMP) | voltage_base, Spikecount |
| | SearchHoldingCurrent | bpo_holding_current |
| | SearchThresholdCurrent | bpo_threshold_current |

Table S1: List of protocols and features used for optimizations for each e-type

### S1 Comparing rate-based contributions to local orientation-selective input

We observed in our model (Figure 11b) and also in the model from Billeh et al., 2020 (Figure S12) that cOSI mostly decreased with the firing rate of a neuron. As neurons with higher firing rates have a larger influence on local inputs to other neurons, this effect would reduce the overall efficacy of local connectivity in enhancing orientation selectivity. Importantly, the strength of this biasing effect is influenced by the tailedness of the firing rate distribution, such that a heavy-tailed distribution gives more influence to the less selective, faster spiking neurons.

As mentioned in the introduction, the topological features of morphologically-constrained connectivity lend themselves to amplifying the activity and robustness of a subset of neurons. This can be seen in our model, which has a strongly skewed firing rate distribution. The model from Billeh et al., 2020 had a firing rate distribution which, while skewed, was much more evenly spread and less tailed (not shown). In fact, the spread in Billeh et al., 2020 was less tailed than in the experiment, despite the sampling biases of the latter.

We considered that the differences in the efficacy of tuned connectivity may be due to differences in the firing rate distribution, but a comparison of the rate-weighted cOSIs normalized by mean cOSI between the two models showed comparable numbers (0.55 and 0.59), and thus cannot explain the difference between the models.

### S2 Additional limitations

7

We assigned orientation and direction selective LGN neuron models only to afferents assigned to originate from the LGd-sh (based on Cruz-Martín et al., 2014 and Bickford et al., 2015) which target superficial layers and L6 in our model. However, other experiments demonstrate that orientation-selective boutons are present in L4 as well (Kondo & Ohki, 2016; Sun et al., 2016). We do not see any reason to believe that this could have influenced our results.

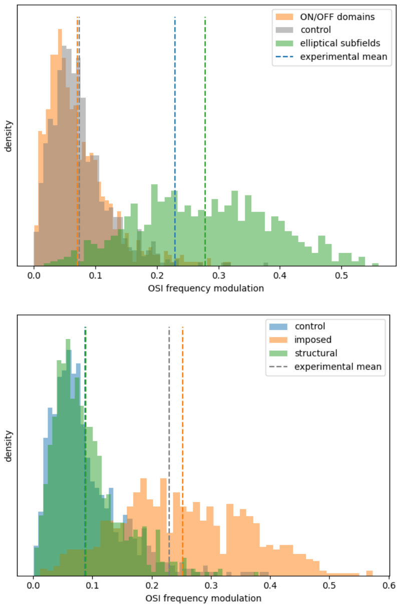


Figure S1: F1-OSI at temporal frequencies of 4Hz and 8Hz

Our method of creating a retinotopic map did not produce the anisotropy observed by some sources (Kalatsky & Stryker, 2003; Schuett et al., 2002), where a displacement along the cortical surface represented a proportionally greater displacement in the horizontal axis of the visual field than the vertical. This was not a deliberate omission, although it did make our analysis a great deal simpler. Had retinotopic anisotropy been present, we would, in basing our connectivity on anatomy, have observed horizontally elongated receptive fields and local connectivity, and we would have needed to devise compensatory mechanisms as in Billeh et al., 2020. In the presence of cortical anisotropy, strong anatomical determination of connectivity may be more difficult to reconcile with orientation tuning.

We mistakenly excluded thick-tufted layer 5 pyramidal neurons from the model (Methods 4.1.1), which have unique apical dendritic morphology. This may have influenced our estimates of connectivity and innervation to layer 5 excitatory neurons. As this cell group is not specifically involved in any of the model failures that form our key results, we do not believe that this has any influence on the strength of our conclusions.

Spontaneous firing rates of Sst-expressing neurons were very low in our simulations. While Ma et al., 2010 also shows low firing rates, they were still significantly higher than in the model. Firing rates during drifting grating stimulation were similar to excitatory firing rates.

### S3 Additional figures

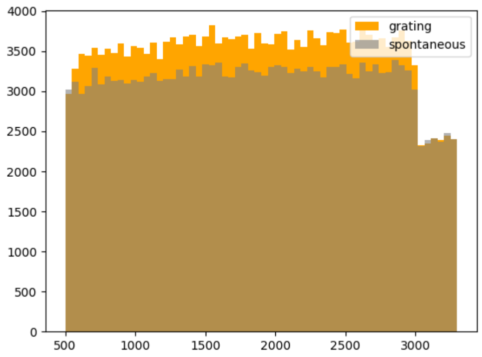


Figure S2: PSTH (grating versus spontaneous) for early model version with facilitating thalamocortical synapses

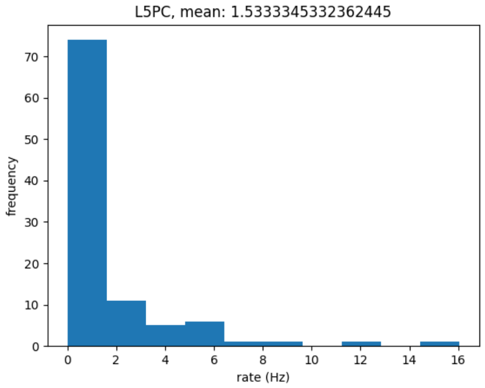


Figure S3: Firing rate distribution for idealized simulations with homogenized emodels

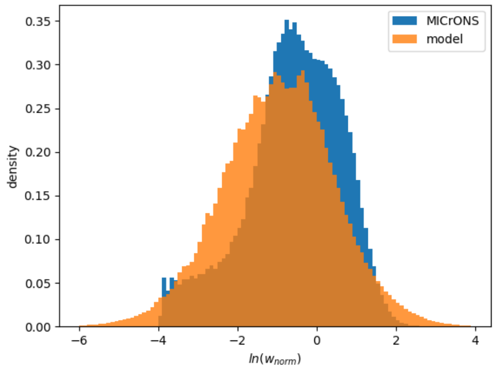


Figure S4: Natural logarithm of connection strength, normalized by mean connection strength in MICrONs data and the first rewired model. The slightly increased spread on the logarithmic scale corresponds to a major increase on a linear scale, with the model having a coefficient of variation of 2.8 and the experiment 1.1

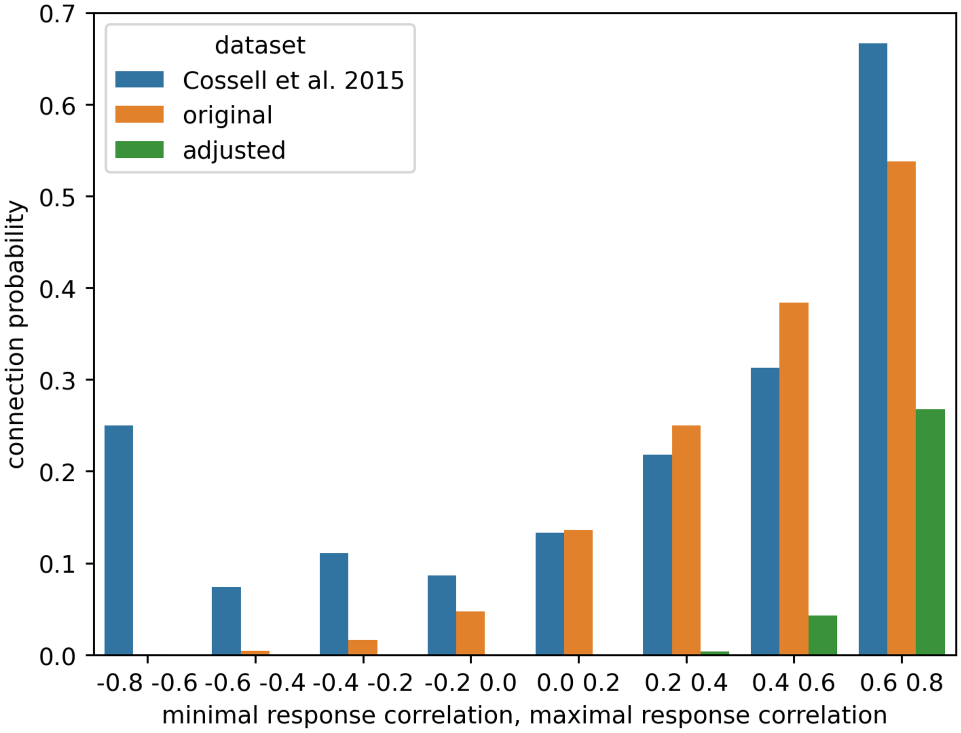


Figure S5: Connection probability of the models compared to Cossell et al., 2015

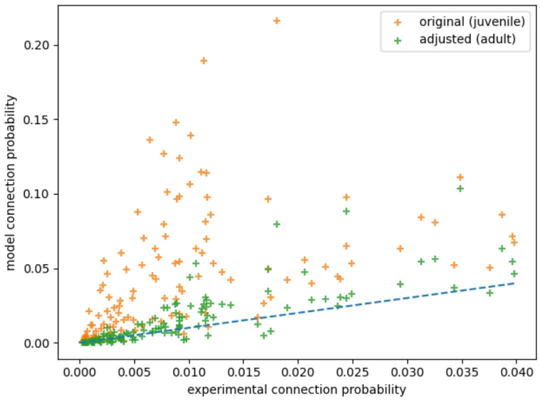


Figure S6: Excitatory connection probability for original and adjusted model compared to MICrONS dataset, layer to layer

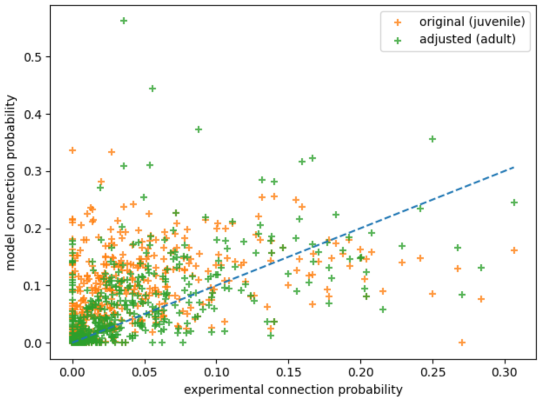


Figure S7: Inhibitory connection probability for original and adjusted model compared to MICrONS dataset, from layer and mclass to layer and mclass

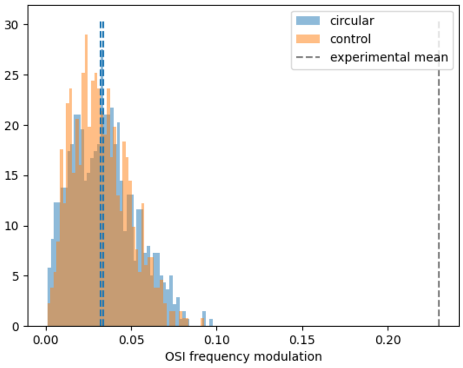


Figure S8: F1-OSI at 2Hz for circular receptive fields with and without segregated ON/OFF domains. We see very little effect from ON/OFF domains

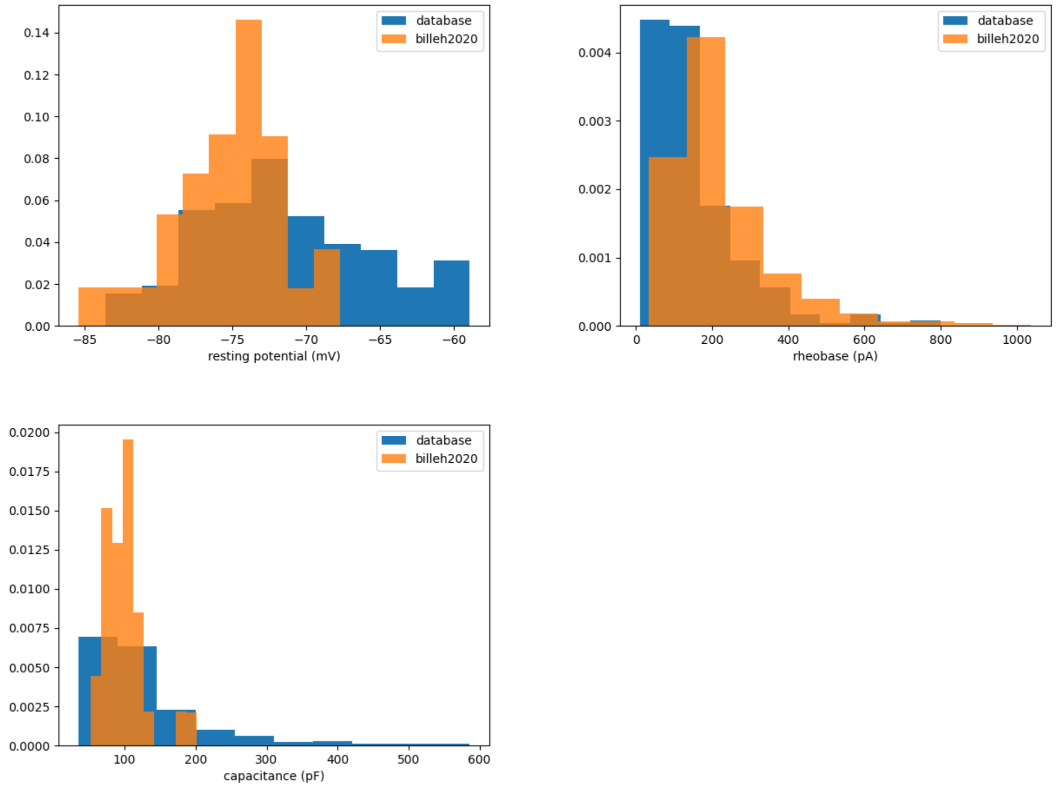


Figure S9: resting potential, threshold current, and capacitance distributions for layer 4 excitatory neuron models in Billeh et al., 2020 (curated cell models), and in our model (sampled directly from the allen cell types database)

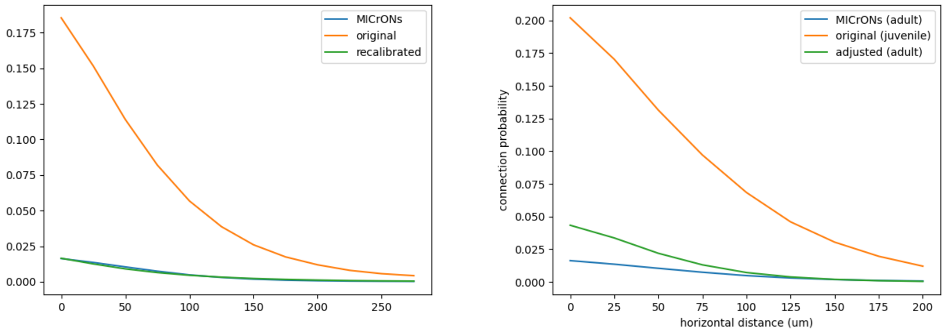


Figure S10: Distance dependent connection probability (excitatory to excitatory) in the original model and recalibrated model compared to Consortium et al., 2021. The upper figure is that predicted by the conditional probability distribution, the lower is the actual connection probabilities in the model itself. Both an increased distance-dependence and greater connection probability are seen in the model, which are absent from the calibration.

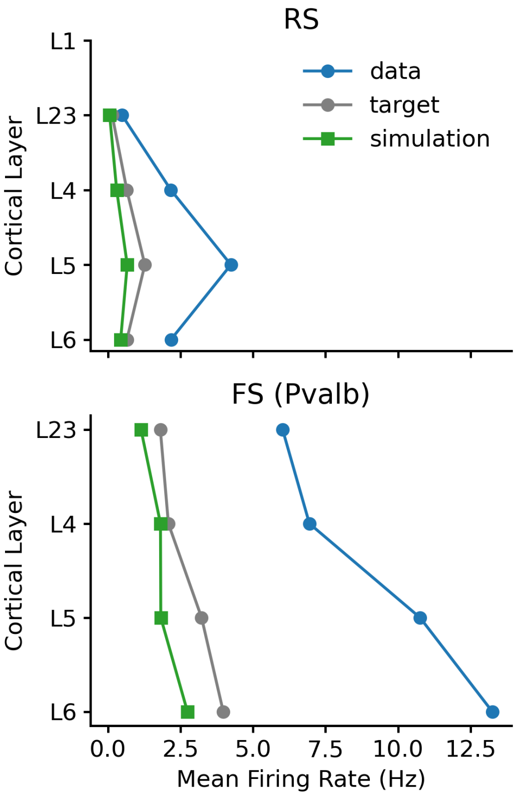


Figure S11: firing rate calibration for thalamocortical-only model

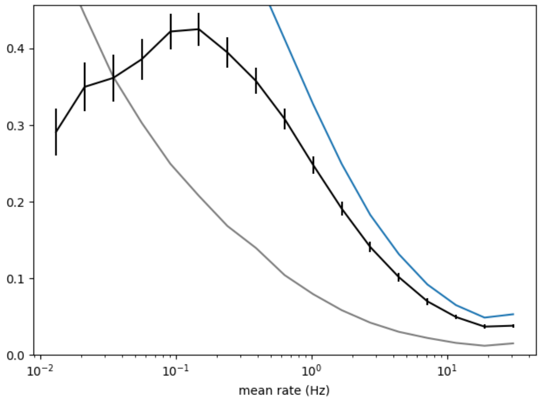


Figure S12: cOSI by mean firing rate for Billeh et al., 2020

### S4 Analysis of alternative explanations for different results

We simulated drifting gratings for a few variations of our model to rule out possible explanations for the discrepancy with other models.

We ruled out some easily testable explanations:

- **insufficient determination of connection strength by orientation preference**: scaling connection strength based on similarity of orientation preferences did not improve OSI (Figure S13b)
- **insufficient phase-dependent connection strength** linearly scaling synapse strength based on phase had little to no effect on OSI (Figure S17)
- **absence of spike-threshold adaptation**: use of neuron models with spike-threshold adaptation from Billeh et al., 2020 did not improve OSI (Figure S14)
- **presence of short-term synaptic plasticity**: abolishing short-term plasticity in the model did not improve OSI (Figure S16)

We additionally checked whether the reduced orientation selectivity compared to experiment could be due to over-estimation of thalamocortical current. Our estimate was based on Lien and Scanziani, 2018, but they note that cortical silencing elevated thalamic firing rates compared to baseline. Baseline firing rates were around 80% of experimental. Lowering thalamic input strength by 20% did not improve OSI S15.

Figure S13

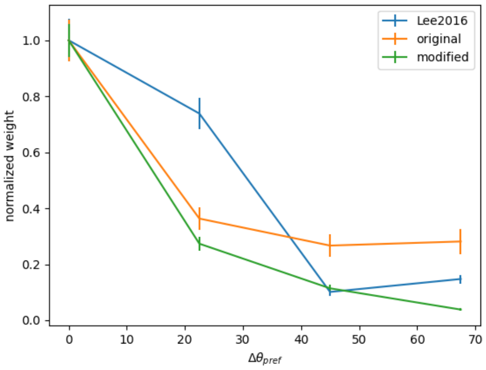


(a) Connection strength scaling to increase correspondence to orientation preference difference. Resulting pearson correlation is 0.1

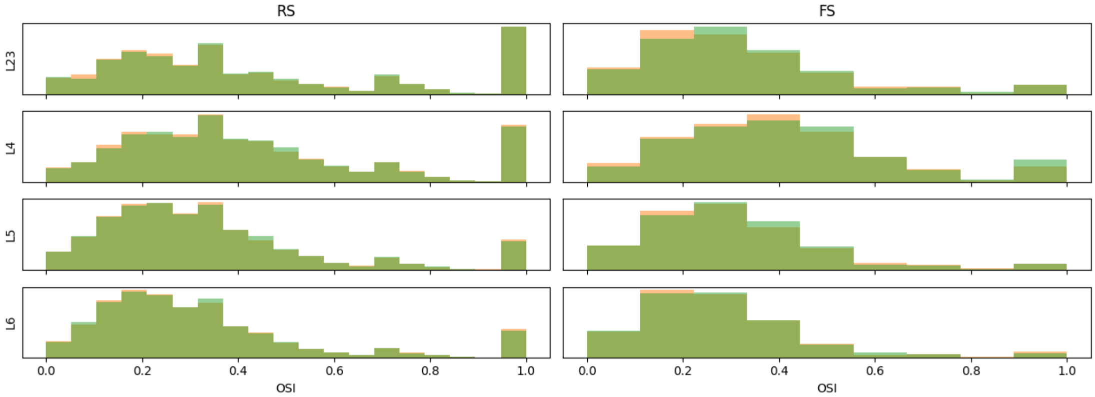


(b) Comparison of orientation selectivity index before (orange) and after (green) connectivity scaling.

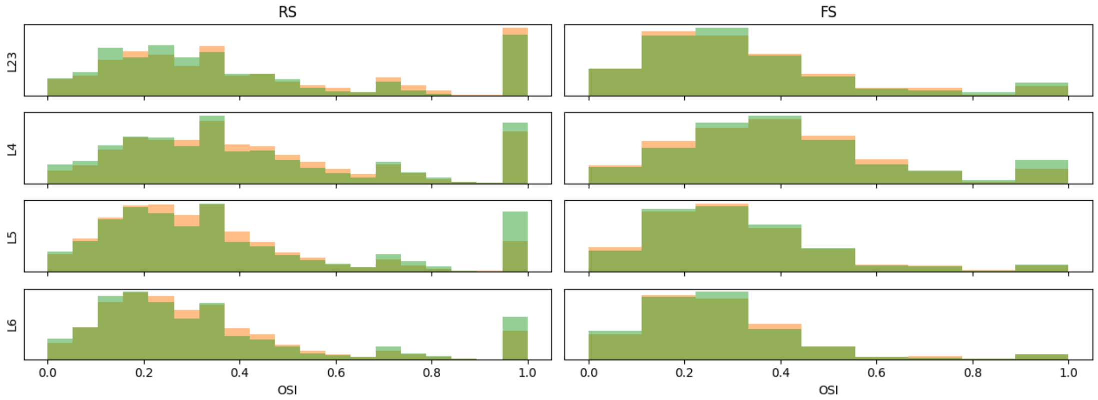


Figure S14: OSI of model with cell models replaced with those from Billeh et al., 2020 (green) compared to the original (orange). Note that orientation selectivity is slightly lower with their cell models.

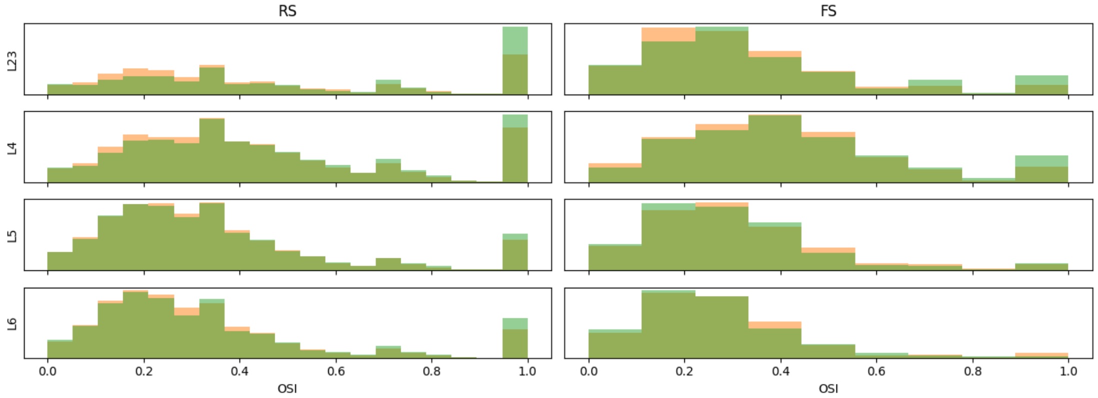


Figure S15: OSI of model with reduced thalamocortical innervation (green) compared to base model (orange).

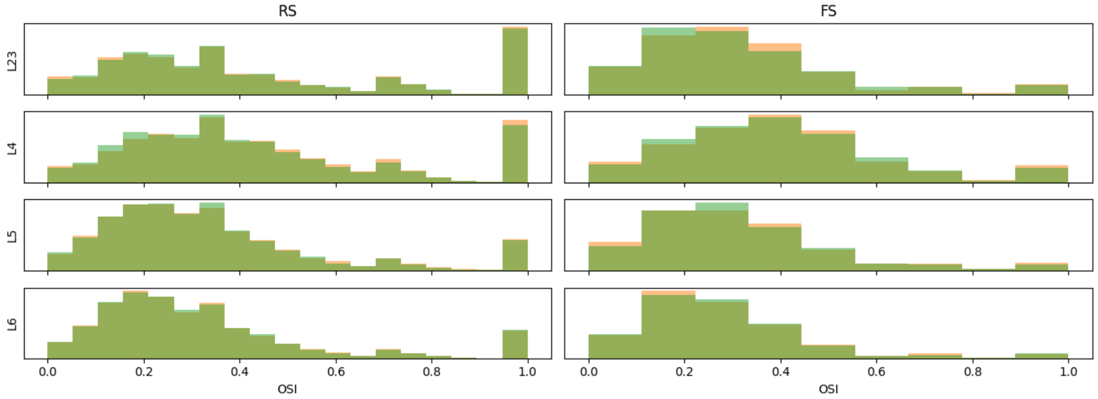


Figure S16: OSI of model without short-term plasticity (green) compared to base model (orange)

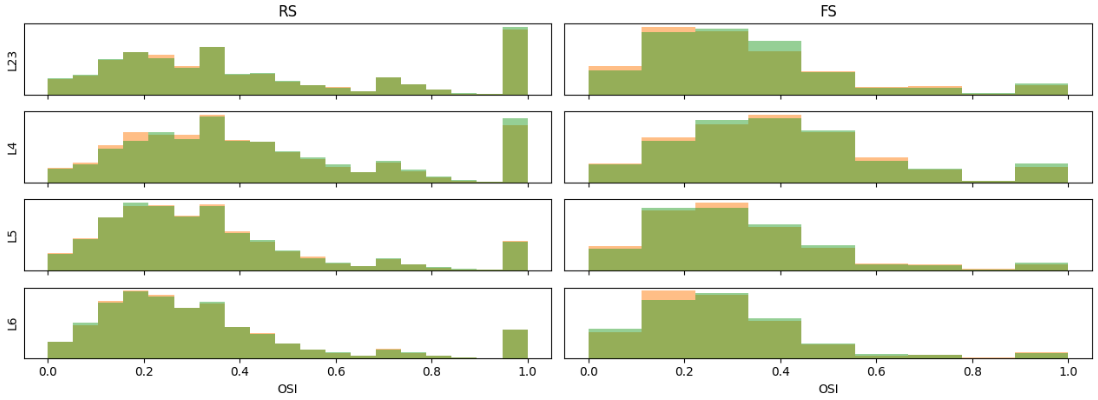


Figure S17: OSI of model with phase-scaled connectivity (green) compared to base model (orange). synapse strength was scaled linearly with the inverse of retinotopic distance, projected in the direction of the target neuron's preferred orientation. The scaling coefficient was set so that connection strength would be 0 when the projected distance is 4 degrees or more. remaining connections were scaled up to preserve mean synapse strength
